# Dissection and reconstruction of the colorectal cancer tumor microenvironment

**DOI:** 10.1101/2023.11.01.565169

**Authors:** Nicolas Broguiere, Luis Francisco Lorenzo Martin, Simone Ragusa, François Rivest, Nathalie Brandenberg, Sylke Hoehnel-Ka, Devanjali Dutta, Marième Gueye, Daniel Alpern, Bart Deplancke, Lana Kandalaft, George Coukos, Krisztian Homicsko, Matthias P. Lutolf

## Abstract

Patient-derived organoids (PDOs) are the reference *in vitro* human disease models. However, the utility of colorectal cancer (CRC) PDOs is hindered by the lack of a tumor microenvironment (TME). To address this limitation, we built a living biobank of CRC PDOs with autologous stromal and immune TME. We characterized the original tumors and traditional monocultures using single-cell RNA-seq (scRNA-seq) and whole exome sequencing (WES) to obtain insights into cell type selection and phenotypic drift in culture. Subsequently, we developed culture conditions supporting all cell types to recapitulate the CRC-TME around PDOs. From the transcriptomes of >180k cells obtained from 260 such co-cultures, we illuminated the mutual influence of cells within CRC tumors. Based on original tumor data, atlases of predicted interactions and transcriptional networks elucidated why monocultures were altered and suggested that TME reconstruction more accurately reflected original tumor behavior. We found that inflammatory signals were absent *in vitro* and recovered upon co-culture with tumor-infiltrating lymphocytes (TILs). We also functionally confirmed that stromal, not cancer cells, mediated immune evasion. Additionally, stroma induced an invasive phenotype in cancer cells. From this deep dive into CRC-TME interactions, we built the human CRC-TME atlas (https://crc-tme.com/), an online portal for interactive exploration of gene expression data, prediction of cell-cell interactions at the pathway and receptor/ligand levels, transcriptional networks, and more. We anticipate PDO cultures with reconstructed TMEs will be valuable for discovery efforts, preclinical studies, and personalized medicine, with the atlas as a framework and inspiration for future CRC-TME studies.

## Introduction

Cancer persists as a significant challenge for medical research due to its inherent heterogeneity. Each cancer subtype is commonly associated with numerous mutations that yield millions of possible combinations, coupled with diverse patient backgrounds, to make each tumor unique. For this reason, personalized medicine approaches are highly sought after. Notably, patient-derived organoids (PDOs) have emerged as a promising platform to study patient-specific cancer biology and drug response in a timely, affordable, and reasonably biologically accurate fashion^1^. A significant limitation to the accuracy and utility of standard PDO cultures is their sole propagation of epithelial cells. Therefore, we envisioned a new generation of PDOs that include a reconstructed tumor microenvironment (TME). The non-malignant cells comprising the TME within tumors are expected to affect the cancer cells’ growth, survival, migration, differentiation, and drug response.

In this study, we established a biobank of human colorectal cancer (CRC) PDOs with their associated stromal and immune TME. We cultured the cancer epithelium (EP), tumor-infiltrating lymphocytes (TILs), and cancer-associated fibroblast-like cells (CAFs) from tissue samples. We also derived antigen-presenting cells (APCs) from autologous peripheral blood monocytes (**Figure 1A**) and co-cultured the PDOs with their TME using optimized protocols.

**Figure 1.**
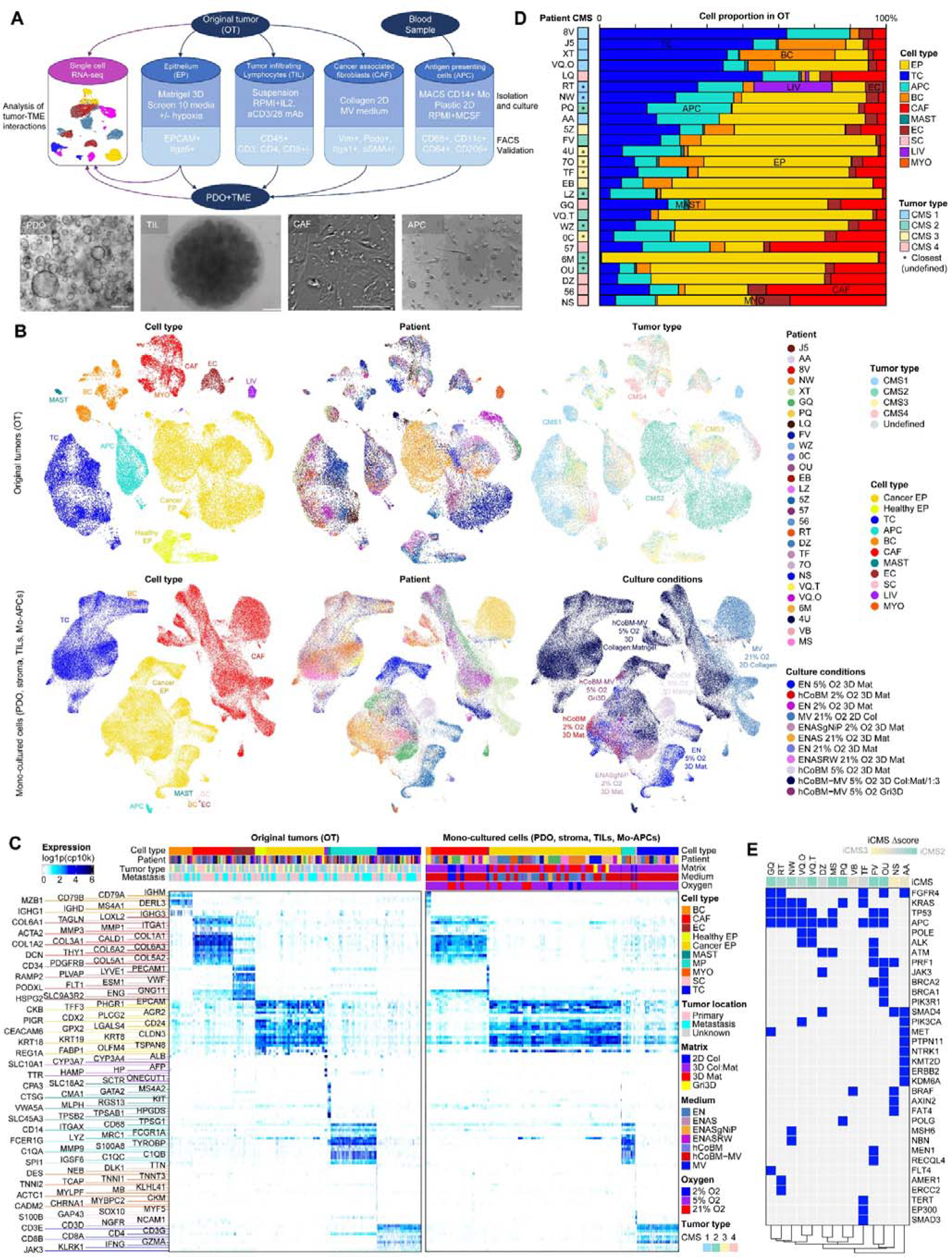
(A) General workflow for generating the living biobank: cell types with their culture conditions, markers, and brightfield view. Scale bars: 100 μm. (B) UMAPs with Scanorama integration show the identities of the cells profiled from OTs or monocultures. (C) Heatmap of cell type markers used to classify the cells into overarching categories, and their expression in vitro and in vivo. (D) Consensus molecular subtypes (CMS) and cell proportions in OTs. (E) Mutations in cancer cells, from WES of PDOs with hierarchical clustering, and annotated with the difference between the iCMS2 and iCMS3 scores in the corresponding PDO lines.

We profiled the original tumors (OTs) and monocultured cells as well as the reconstructed PDO+TME to generate a multimodal atlas using single-cell RNA-seq (scRNA-seq). We complemented the single-cell data with whole exome sequencing (WES) of the cancer cells, fluorescence-activated cell sorting (FACS) of monocultured cells, and functional data on cell cultures. The resulting atlas of CRC-TME interactions highlights common trends and patient heterogeneity. With this living biobank, we gained an opportunity to identify critical interactions between cell components. We made the data explorable on an online portal, which we called the CRC-TME atlas. The CRC-TME atlas features web applications for interactive exploration of the various cell types in CRC tumors, their gene expression, cell-cell interactions, as well as similarities and differences between *in vitro* and *in vivo* single-cell profiles for both monocultured and co-cultured cells.

To meet the software requirements for building the atlas, we developed a suite of tools in the R programming language designed specifically for the analysis of complex tumor samples. The main features include single-cell level classifiers based on signature gating and single-cell level variant/mutation analysis, emphasizing documentation and reusability for other challenging single-cell projects. The package, named “Bundle of Useful scRNA-seq Gating Tools Extending seuRat”, or “burgertools”, is available at github.com/nbroguiere/burgertools.

## Results and discussion

### Profiling of the original tumors and establishment of the CRC+TME living biobank

We dissociated fresh surgery or biopsy-derived OT samples to single cells. We obtained cell quantities on the order of 100 k from needle-core biopsies and 10 M from surgery samples, with strong variability. We immediately used a fraction for scRNA-seq profiling to generate a reference of the in vivo phenotype. We established cultures with the remaining cells and froze any excess.

The various cell types in CRC tumors rely on drastically different growth factors and extracellular matrices (ECMs), which we leveraged to expand various populations selectively (**Figure 1A**). This selective growth strategy from a heterogeneous mixture minimizes cell stress and yields mostly pure populations, as confirmed by flow cytometry/immunocytochemistry (**Figure S1**).

We expanded the cancer cells in 3D using basement membrane extract (Matrigel) and patient-specific media selected from an initial screen (**Figure S2**). Conditions ranged from no growth factor to EGF, Noggin, TGFßi, p38i, gastrin, Nicotinamide, Prostaglandin-E2, R-Spondin-1, and Wnt, henceforth referred to as ENASgNiPRW. We tested hypoxia vs. normoxia and found that the cultures functioned equally well or better under the former while retaining physiological relevance. Therefore, we cultured PDOs under 5% O_2_ for all further experiments.

We grew stromal cells on stiff collagen and found the best growth in a rich medium optimized for microvascular cells, namely EGM2-MV. All the factors in this medium benefited the CAFs (**Figure S3**). We grew TILs in suspension culture using the traditional immune cell medium, RPMI+10%FBS, with the key growth factor IL2. While APCs are an essential part of the tumor-associated innate immune system, they cannot be expanded from the tumor itself. To address this limitation, we isolated CD14^+^ monocytes from blood samples collected simultaneously with tumor samples and differentiated them *in situ*. We then profiled these monocultured PDOs/primary cells with scRNA-seq and established a baseline of cell phenotype in standard cultures.

We processed samples from 31 patients and successfully derived OT scRNA-seq for 26, totaling 35k cells (**Figure 1B**). 23 samples were from surgery and had a PDO establishment success rate of >95%. However, several PDO lines grew too slowly to have utility for experiments. 8 samples were from needle-core biopsy and had a PDO establishment rate of only 37.5%, which underscored the dependence of successful PDO generation on initial cell numbers. We isolated monocytes with high viability from 15 patient blood donations. We attempted CAF cultures for 19 donors, with a ∼95% success rate, and established TIL lines for 18 patients, which were 100% successful. In total, we profiled 65 monocultures with 110k cells across diverse conditions.

We classified cells into the overarching main categories: EP, CAF-like (e.g. CAFs, fibroblasts, myofibroblasts, and pericytes), APC, T cell-like (TC), B cell-like (BC), and endothelial cells (EC), as well as some rare types including MAST-like cells (MAST), myoblasts/myocytes (MYO), liver cells (hepatocytes and cholangiocytes, denoted as LIV), and Schwann cells (SC). Since small populations can be missed in clustering, we developed a custom workflow based on signature gating at the single-cell level, analogous to FACS (**Figure S4A**). The cell type-specific markers used to define these categories were highly conserved across patients and between OTs and cultures (**Figure 1C**). We performed quality control (QC) by cell category from these clustering-free classifications (**Figure S4B**), which was essential for these highly heterogeneous cancer samples. To distinguish cancer cells from residual healthy epithelium, we developed a gating approach based on single-cell mutation/variant data (**Figure S4C**). We confirmed the robustness of the approach by cross-checking known cancer mutations in single cells (**Figure S4D**). The suite of tools that we developed to perform these gating, QC, and single-cell variant analyses are available in burgertools.

Furthermore, CRCs are commonly stratified based on their consensus molecular subtype (CMS)^2^, which we obtained from pseudo-bulk aggregated RNA-seq data (**Figure 1D**). The CMS subtype is mainly determined by immune cell content (contribution towards CMS1) vs. CAF content (contribution towards CMS4). The cancer cells themselves contributed towards either CMS2 or CMS3, depending on their metabolic profile (**Figure S4E**), which can be seen as a cancer-intrinsic CMS^3^. Patients from all CRC subtypes were well represented among our donors.

We obtained mutations in cancer cells for 18 patients using WES of PDOs (**Figure 1E**). All common CRC-associated mutations were represented in the biobank. As expected, mutations correlated with the CMS scored on pseudo-bulk of cancer cells only, referred to as intrinsic CMS (iCMS): typical *APC/KRAS/TP53* mutants and closely related profiles scored high on iCMS2, whereas *BRAF* mutants and an atypical patient scored higher on iCMS3. The main mutated pathways are those typically associated with stem cell maintenance in the intestinal epithelium and organoids. Notably, Wnt, EGF, and other RTK binding growth factors, such as HGF/FGF/IGF, have activating mutations. Conversely, BMP/TGFß and DNA repair and inflammation-related pathways acquire loss of function mutations. The iCMS correlated between *in vitro* and *in vivo* (**Figure S4F**).

In low dimensional embeddings (**Figure 1B**), we found that combined datasets clustered primarily by cell type, secondarily by patient, and thirdly by CMS *in vivo*, or culture condition *in vitro*. These findings qualitatively hinted at sources of heterogeneity in and across CRC tumors/cultures and their relative importance.

### PDOs significantly reproduce patient heterogeneity in cancer cells

The cancer cells were heterogeneous across patients, with unique combinations of intestinal stem and differentiated cell markers, as well as various epithelial-to-mesenchymal transition (EMT) related genes. To quantify the heterogeneity conserved in the PDOs, we computed patient-specific markers in vitro and in vivo. Notably, PDOs shared many markers with their OTs and few with other tumors (**Figure 2A**). Many markers were relevant intestinal epithelium or cancer genes, such as *SOX2/OLFM4* (stem), *CCK/CHGB* (enteroendocrine), *DEFB1* (Paneth), or *MMP7* (EMT) (**Figure 2B**). We aligned cancer cells in all OTs vs. PDOs based on the union of these markers (**Figure 2C**) and found that all PDOs clustered with their OTs. Additionally, the Spearman correlation of the patient-specific markers expression was significantly higher for matched OTs and PDOs (**Figure 2D**), and not affected by the culture conditions (**Figure 2E**).

**Figure 2.**
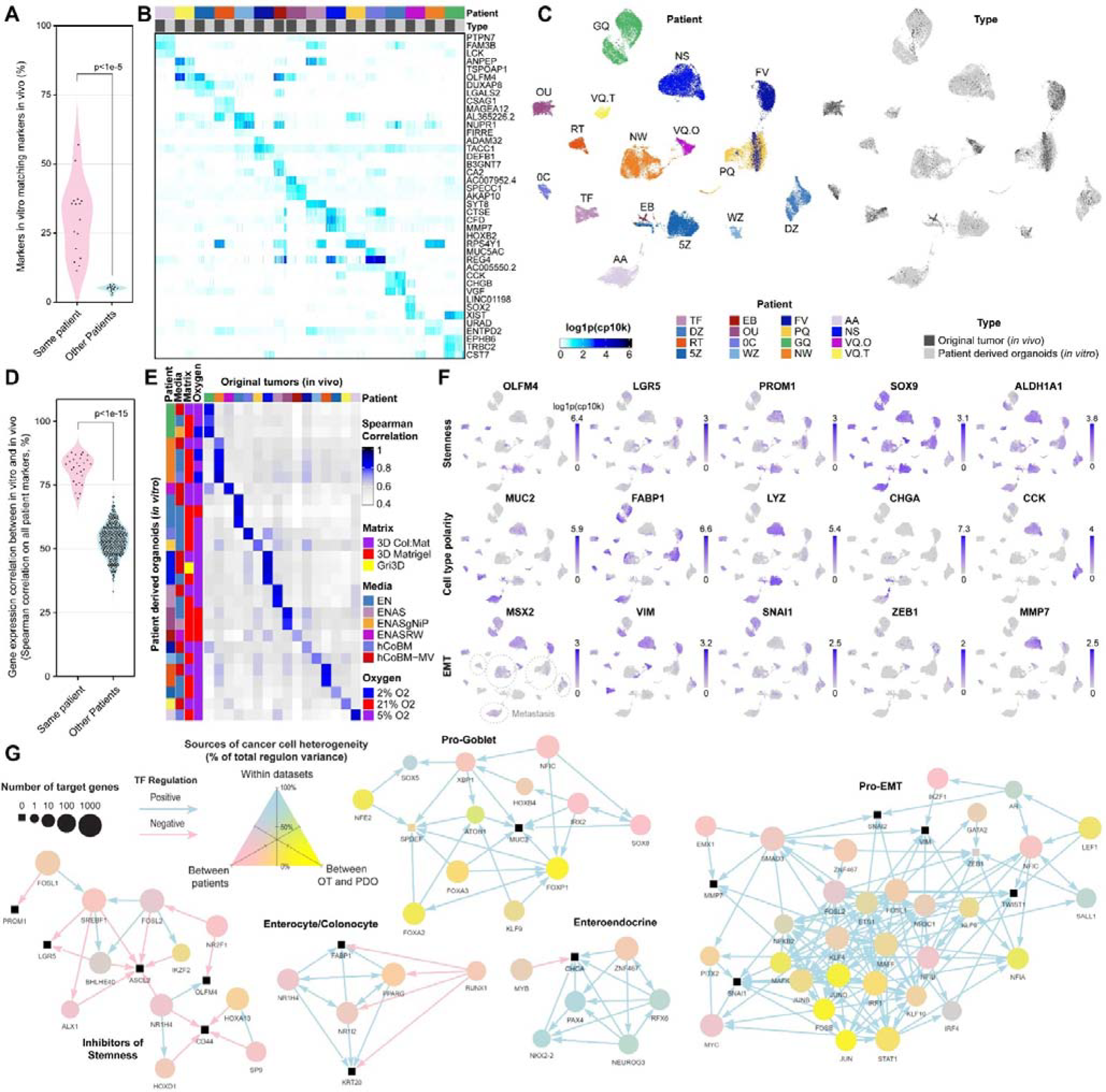
(A) Percentage of patient-specific markers that match between in vitro and in vivo, and two-sided t-test. (B) Heatmap of the top 3 conserved patient-specific markers for all patients, for cancer cells profiled both in vitro and in vivo, with OTs and PDOs shown side-by-side. (C) UMAP of cancer cells in vitro and in vivo, computed based on the union of all conserved patient-specific markers and harmony-aligned. (D) Spearman correlation between the expression in vitro and in vivo, based on the union of conserved patient-specific markers, and two-sided t-test. (E) Heatmap of the Spearman correlations in D, organized by patient and culture condition. The patient color scale from C applies. (F) Expression of key markers of stemness, polarity, and EMT in cancer cells, on an overlay of OT and PDOs. Cells originating from metastasis are highlighted (bottom left subpanel). (G) Transcriptional networks in epithelial cells as predicted by SCENIC. Each subnetwork highlights the TFs regulating key genes (black squares). Three networks are further restricted to only positive or negative regulators. Colors indicate the origin of the variance of the regulons scores, obtained by average log expression of the regulons target genes.

We found only limited congruencies between various stem or EMT markers, and the intestinal cell type differentiation markers often recombined unpredictably, which hindered our assignment of clear “stem-like” or “goblet-like” phenotypes (**Figure 2F**). This seemingly random co-expression of intestinal markers likely reflects the high dysregulation of cancer cells despite of their close relationship to the epithelium of origin. However, several patients had a clear enteroendocrine differentiation axis, and over half of the patients had some enterocyte-like polarity gradient. These partial differentiations did not necessarily reduce stem and progenitor gene expressions (e.g., *SOX9* was highly expressed in almost all cancer cells).

Cancer stem cells (CSCs) have often been defined based on one or two surface markers used in FACS sorting^4^. The caveat is that the cells isolated and defined as CSCs vary depending on the chosen marker(s) as most cancer cells are positive for at least one CSC marker. Surprisingly, the cancer cells biopsied from metastases exhibited low canonical EMT marker expression, suggesting re-epithelialization after migration, typically to the liver, or invasion without EMT.

To better understand the sources of variation between and within patients, we predicted the transcriptional networks within epithelial cells using the SCENIC workflow. The resulting transcriptional network has 182 positive and 186 negative regulons, with 57 k edges, including 3.3 k between transcription factors (TFs), which is difficult to visualize at once. The CRC-TME web portal offers various filtering and visualization options to conveniently explore interesting sub-networks.

*Ab initio* prediction of TF targets was remarkably consistent with current experimental knowledge (**Figure 2G**). For example, the predicted regulators of the enteroendocrine marker CHGA, with strong intratumor heterogeneity, included the canonical enteroendocrine TFs PAX4, NKX2-2, NEUROG3, and lesser known RFX6^5^. These TFs reinforced each other, a classical mechanism for identity maintenance^6^. Goblet markers are partially regulated by well-known TFs such as ATOH1, XBP1, and FOXA2/3, which vary between patients and *in vitro* vs. *in vivo*. The TFs NR1H4 and NR1I2 (bile acid response) and PPARG (lipid response) governed enterocyte polarity, marked by *FABP1* and *KRT20*. Again, these TFs reinforced each other and were antagonized by the progenitor TF RUNX1. Additionally, FOSL1/2 and the lipid master controller SREBF1 inhibited stemness, and the primary source of variation was patient diversity. FOSL2 was a key predicted positive regulator of EMT, together with SMAD3 (response to TGFß family), which suggests opposing states between stem/progenitor and EMT. This effect is observed in response to TGFß, where EMT is induced at the expense of cell proliferation, which varied predominantly between patients. Other regulators of EMT included AP-1 and inflammatory response factors, such as STAT1, and were markedly different between *in vitro* and *in vivo*. This marked difference highlights the limitations of and need for improvement of *in vitro* models.

### Stromal cell diversity in OTs and CAF cultures

Stromal cells exhibited remarkable diversity in OTs and consisted predominantly of stromal cells and ECs (**Figure 3A**). Based on the expression of canonical markers (**Figure 3B**), we further categorized ECs into lymph, sinusoidal, and vascular types, where the latter comprised the majority and was the most proliferative. Schwann cells were uncommon but present in several tumors (**Figure 3C**). Intriguingly, the Schwann cells expressed genes usually associated with APCs – a phenotype previously described as an inflammatory response in nerves ^7,8^. Myoblasts/myocytes were only present in one tumor, likely from sampling submucosal muscle infiltration.

**Figure 3.**
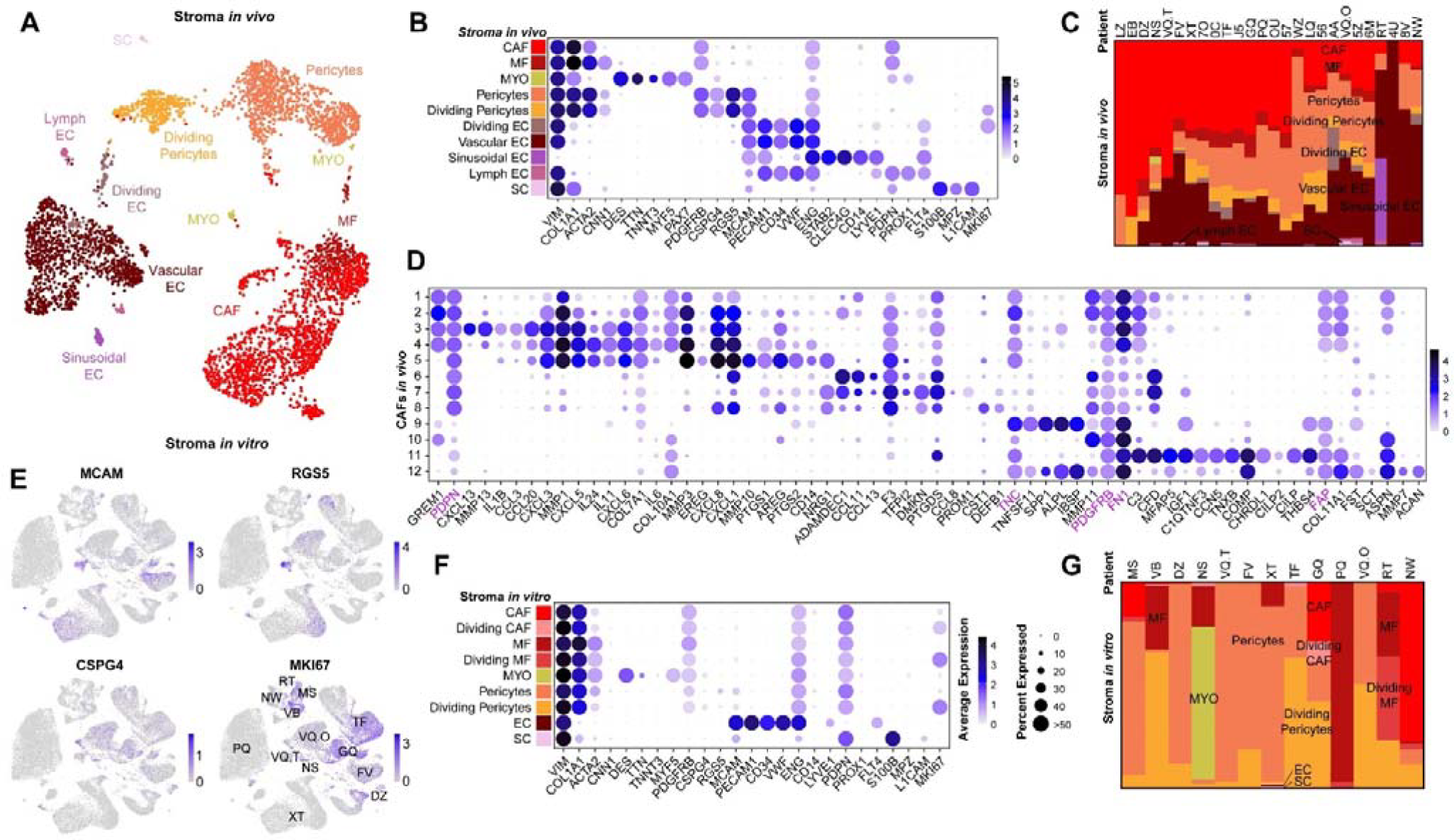
(A) UMAP of stromal cells from all OTs (Scanorama integrated). (B) Expression of marker genes by cell subtype for stromal cells in vivo, in which cells categorized as CAF or EC in Fig. 1 are subdivided into finer categories. Stromal cell subtype proportions in vivo per patient. (D) Further subdivision of CAFs into 12 clusters and their associated markers. Canonical-activated fibroblast markers are highlighted in purple. (E) Expression of residual pericyte markers, and a proliferation marker, in stromal/CAF cultures, displayed on UMAP without alignment. The distinct clusters correspond to different patients, annotated in overlay on the MKI67 subpanel. (F) Expression of stromal subtype marker genes in vitro. (G) Proportions of cell subtypes in vitro, dominated by dedifferentiated pericytes.

In most cases, CAF-like cells (*COL1A1*^+^ *S100B*^neg^) constituted the bulk of the stroma. The ACTA2^+^ cells were identified as myofibroblasts (MF), and with additional *CSPG4*^*+*^ *RGS5*^*+*^ *MCAM*^+^ as pericytes. The CAF-like cells that did not fall into these sub-categories were extremely heterogeneous within and across patients. We collected them into 12 groups for illustrative purposes (**Figure 3D**), but found further diversity with finer clustering. Strikingly, the defining markers for activated fibroblasts, such as *PDPN, TNC, PDGFRB, FN1*, and even *FAP*, were heterogeneously expressed among these populations. They were also not necessarily correlated with each other, which implied that the diversity of the phenotypes adopted by CAFs was higher than anticipated. These populations also expressed very diverse growth factors, cytokines, matrix proteins, and matrix remodeling enzymes. Strikingly, we found that pericytes were the only stromal cells that were proliferating and formed a continuum towards MF and CAFs. This suggests that most of the CAFs in CRC tumors differentiate from pericytes.

*In vitro*, the CAF cultures exhibited high heterogeneity across patients, culture conditions, and within a given culture flask. Dozens of CAF populations with distinct cytokine/matrix markers could be distinguished, but most of these cells had residual expression of one or several pericyte markers (**Figure 3E, F, G**). This is consistent with the previous observation that pericytes are proliferative cells that presumably give rise to the various fibroblast populations.

Overall, CAFs seem exquisitely sensitive to their environment both *in vivo* and *in vitro*, which engenders tremendous diversity and rapid phenotypic drift. Nevertheless, deriving from well-defined pericyte precursors might open avenues to study their generation and differentiation, as well as control their polarity with bioengineering approaches, as exemplified with stem cells^9^.

### Immune cell diversity in OTs and TIL cultures

We observed a tremendous diversity of immune cells in vivo, and the cell types and polarity markers we identified showcased the immune reaction to the tumors. The most numerous, diverse, and proliferative cells were the T-like lymphocytes, particularly CD4^+^ T cells. Their main polarities were T_h_ 1, T_h_ 17, and T_fh_ (**Figure 4A, B, C**), which indicated a type I immune response to the tumor and the presence of tertiary lymphoid structures. These cells also highly expressed the associated cytokines *IFNG, IL22, IL21, IL17A/F, IL26*, and *CXCL13*. T_h_2 was absent and their associated cytokines *IL4* and *IL10* not expressed in T_h_. Two of the three T_h_1 clusters were cytotoxic-like (CTL), with high expression of GZMB and other cytotoxic markers. CD4^+^ KLF2^high^ CCR7^high^ naive T_h_ were abundant but did not strongly express cytokines: these cells are putatively newly recruited and acting as a reserve for activation and differentiation.

**Figure 4.**
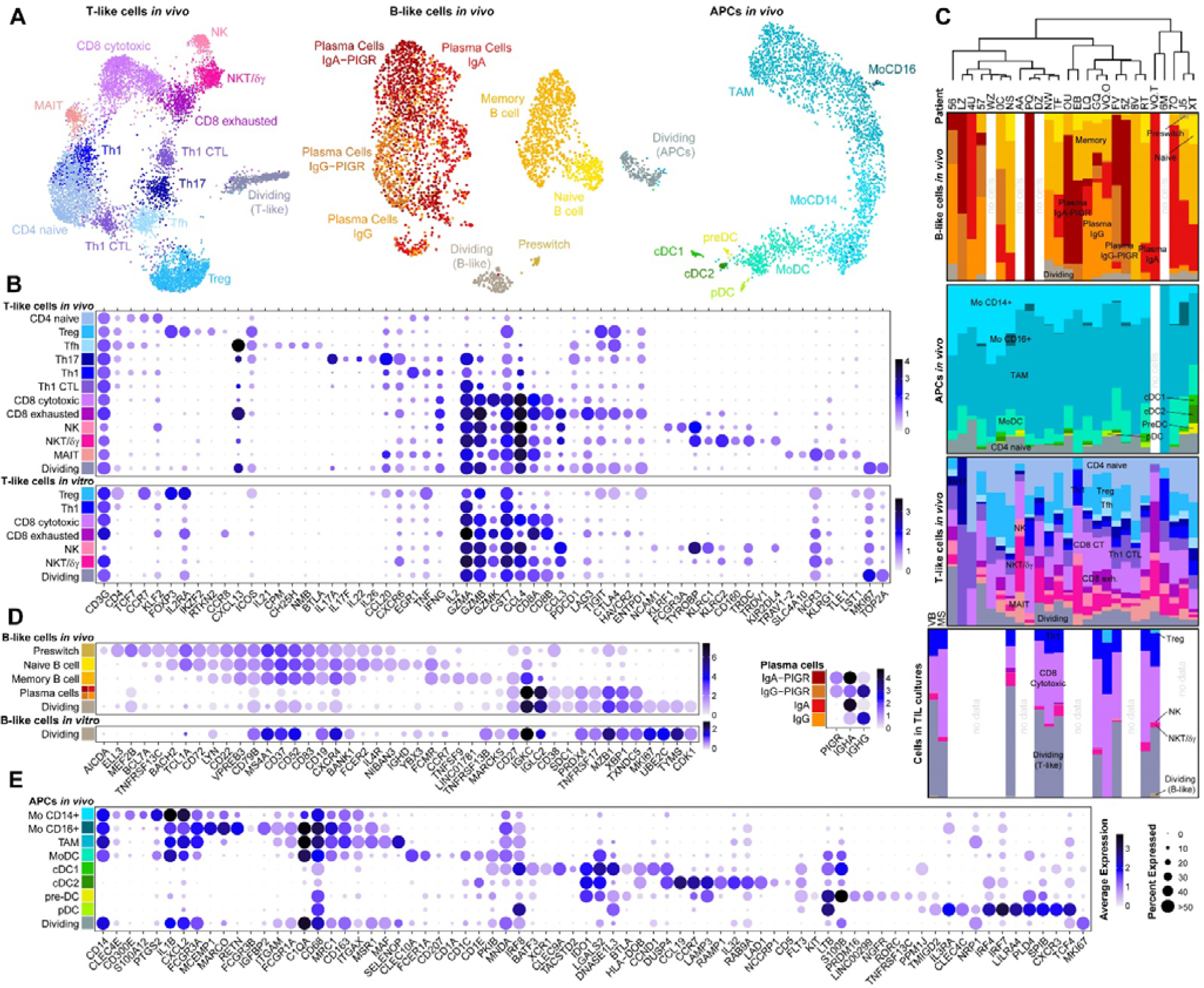
(A) UMAPs showing subtypes among APCs, T-like and B-like cells in original tumors, harmony aligned except for B-like. (B) T-like cell subtype markers in OTs and TIL cultures. (C) Cell subtype proportions in OTs and TIL cultures. (D) B-like cell subtype markers in OTs and TIL cultures. (E) APC cell subtype markers in OTs.

The next most abundant T-like population was CD8^+^ cytotoxic T cells. Both CD8^+^ and CD4^+^ CTL had high expression of the exhaustion markers *PDCD1 (PD1*), *LAG3, TIGIT*, and *CTLA4*, as well as *HAVCR2* (CD39). This expression pattern suggests that tumor survival and immune escape is likely due to T cell exhaustion. T_reg_ were abundant and strongly expressed *TIGIT* and *CTLA4*, as well as *IL10* and *HAVCR2*, which might further contribute to cytotoxic T cell exhaustion. Their abundance might be explained by the high expression of *CCL20* by T_h_17, a cytokine previously attributed to myeloid and cancer cells, and known to recruit inhibitory myeloid-derived suppressor cells and T_regs_^10^.

Rarer lymphocytes included mucosa-associated innate lymphocytes (MAIT). The MAITs expressed cytokines such as CCL4, which is responsible for immune cell recruitment to the tumor, and TNF. We also observed NK, NKT, and γδ-T cells. NK cells were predominantly *NCAM1*(CD56)^dim^*CD16*^+^ and included *KLRF1*^+^ mature and *FGFBP2*^+^ activated populations. Interestingly, the NKT cells expressed the γδ-T cell receptor and the innate lymphocyte transcription factor *PLZF*, which matched a rare population described previously^11^. While these cells are considered promising for immunotherapies^10^, their main proposed mode of action (expression of *IL17A/F*) was not corroborated by our data. Instead, they demonstrated high expression of cytotoxic markers and low expression of exhaustion markers. These innate-like lymphocyte populations might play a key role in activating an initial response to the tumor by acting as a first line of defense and recruiting other immune cells.

We observed a reduction of both diversity and differentiation of the T-like populations in TIL cultures (**Figure 4C**). Naïve TCs, MAIT, T_fh_, T_reg_, were almost entirely lost, with reduced NKT/γδ T cells. Helper T cells reverted to a more generic T_h_ 1 phenotype with largely reduced expression of cytokines (**Figure 4B**). *CD8*^+^ cytotoxic T cells mostly lost the expression of exhaustion markers, which suggested an inhibitory environment in vivo and exhibited much greater proliferation.

The other lymphocytes (∼25%) in tumors were B-like (**Figure 4A, C, D**), primarily differentiated plasma cells (63%), which reflected a mucosal (IgA^+^ PIGR^+^) or systemic (IgG^+^) immune response. The other main types of B-like cells were *CD27*^*+*^*TNFRSF13B*(CD20)^+^ memory (25%) and IgD^high^*CD27*^neg^ naïve (6%) B cells. Memory B cells expressed *TNFSF9* (4-1BB-Ligand), which was consistent with a role in tertiary lymphoid structures, as were pre-switch and proliferating B cells. We observed very few BCs in vitro as the culture protocol was designed for T cells. We envision that future TME reconstruction culture protocols could be extended to BCs.

Finally, APCs were in slightly greater abundance than BCs in OTs and mainly on a differentiation axis from *CD14*^*+*^ monocytes to *MSR1*^+^*MAF*^+^*CD68*^high^*MRC1*^high^*CD163*^high^*ITGAX*^high^ tumor-associated macrophages (TAMs, **Figure 4A, C, E**). A few *FCGR3A*(CD16)^high^ alternative monocytes were also present. Subtype definition was hindered by co-segregation of macrophage polarity markers inconsistent with the literature^12,13^. We also observed that a significant fraction of these cells was proliferating in the tumor. Dendritic cells (DCs) were few and consisted mainly of monocyte-derived dendritic cells (MoDCs), with smaller populations of classical DCs, plasmacytoid DCs (pDCs), and pre-DCs. While marker co-expression did not exactly replicate literature reports^14,15^, they were sufficiently aligned for us to make non-ambiguous assignments to known subtypes. Overall, APC polarizations are more diverse than current reports suggest, and their classification could benefit from meta-studies of single-cell high-dimensional datasets generated in high volume in recent years.

### Monocultured cells lack interactions with other cell types in tumors and have altered gene expression

With simultaneous characterization of cancer and TME cells *in vitro* and *in vivo*, we can observe alterations in tumor cells in the absence of their microenvironment. We can derive information on cancer-TME interactions with potential clinical applications and improve current *in vitro* models.

Toward this goal, we examined differentially activated pathways by cell type based on scRNA-seq data (**Figure 5A**). Subsequently, we linked most differences to four critical aspects of cancer cells. First, we noted the absence of inflammatory signals *in vitro*, such as responses to cytokines like TNF, interferons, and interleukins. Second, proliferation-related pathways significantly increased. Third, we observed altered metabolism, with increased anabolism, particularly for lipid synthesis. Finally, PDOs had reduced EMT-like signatures, with heavy alterations in functions such as organization of the ECM and cell adhesion in a subset of patients.

**Figure 5.**
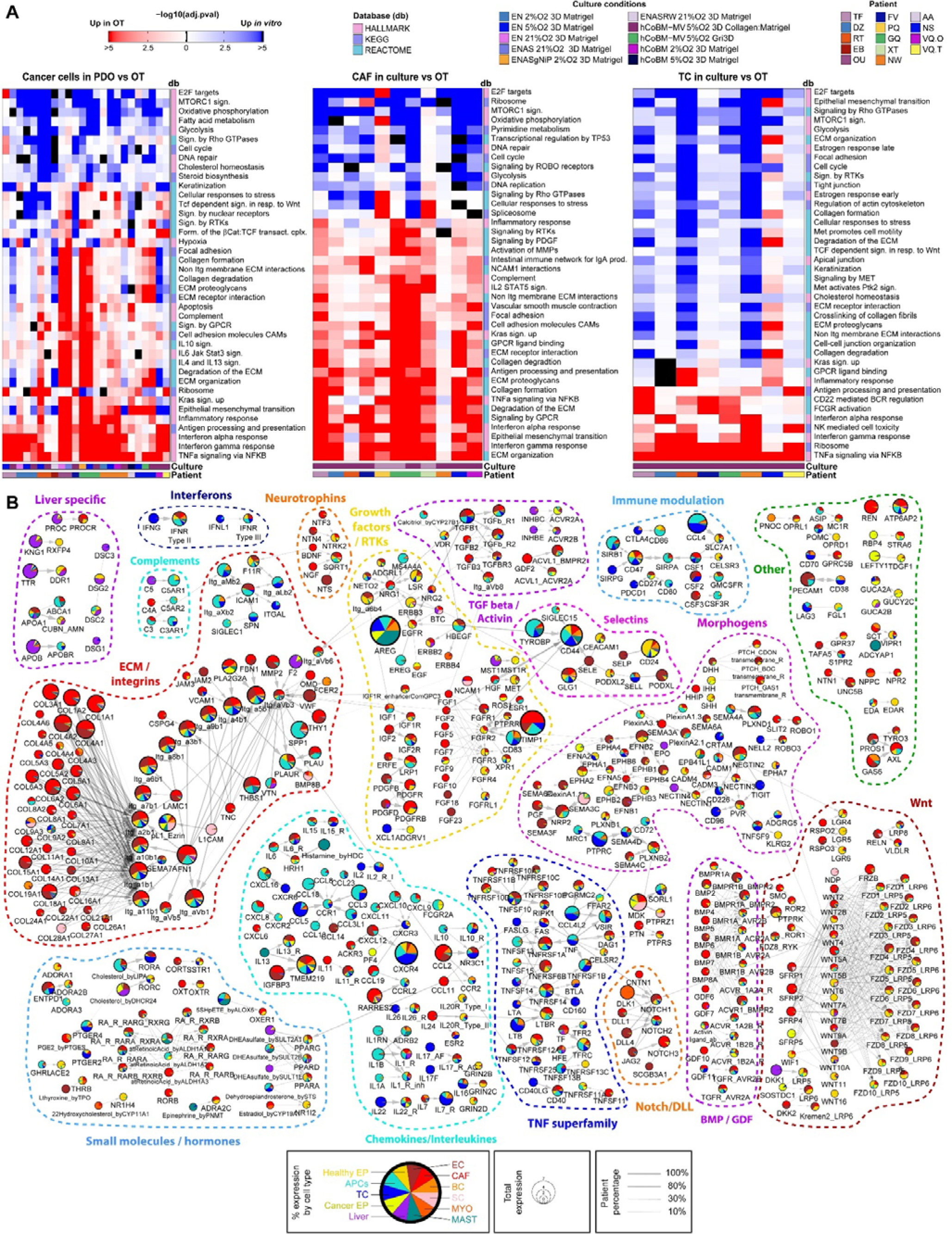
(A) Enriched pathways obtained with fgsea^16^, by cell type comparing cultured cells vs. OTs. Pathways scored are a subset of MSigDB^17,18^ Hallmark, KEGG^19^ and Reactome^20^ – removing duplicates across databases and signatures not relevant to CRC.. (B) Receptor-ligand interactions received by cancer cells, present in OTs but absent in PDOs, as predicted from cellphoneDBv4 and the OT scRNA-seq. Total expression refers to the sum of log-expression across cell types, and the % expression is the ratio of the log-expression of one cell type divided by this sum.

These alterations have significant implications. The efficacy of drugs targeting proliferation could be misjudged in personalized medicine applications based on current PDOs. Another critical study target is EMT, which is not well recapitulated, and the loss of the inflammatory phenotype could greatly impact new immunotherapy studies. Strikingly, CRC cells retained interferon responses and antigen presentation for all the donors in this study, which is promising for T cell targeting strategies (immune checkpoint inhibitors, bispecific and cellular therapy such as TIL and CAR-T). Additionally, stromal cells shifted similarly to cancer cells. We noted a lack of inflammatory signals and increased proliferation *in vitro*, with altered matrix organization and cell adhesion pathways. We nevertheless did not observe marked increases in lipid anabolism. Altered pathways were related to cell identity, consistent with the drastic phenotypic changes described above.

The TC analysis also showed a lack of *in vitro* inflammatory response and increased proliferation at the expense of activity, as expected if inflammation arises from immune/cancer interactions. Unlike PDOs and CAFs, they demonstrated upregulated migration pathways *in vitro*, overall reducing cytotoxic behavior in favor of patrolling.

To find the source of the phenotypic drift, we looked for ligand-receptor interactions absent from the monocultures, using cellphoneDB^21^. From the CRC-TME portal, we can interactively explore the 1.6 k predicted interactions within OTs, encompassing 1 k receptors/ligands in 11 cell types and across 26 patients. We focused on interactions that cancer cells are expected to be missing in PDOs compared to OTs (i.e., interactions where only the receptor is expressed by the cancer cells) (**Figure 5B**). Consistent with previous results, we found that inflammatory cytokines (e.g., IFNγ/λ, chemokines, interleukins, TNF superfamily, complements) and immuno-modulators were expressed by TCs and APCs and absent in PDOs. Furthermore, CAFs contributed diverse ECM components and ENRW analogs, in addition to numerous TGFß/BMP superfamily members, expected to contribute to cell polarization and EMT. Morphogens such as ephrins, semaphorins, nectins, ROBOs, and hedgehog family ligands were abundant in the TME. These morphogens are often examined in development/guidance and could motivate future studies.

Similarly, many hormones are absent and seldom considered for PDOs. For example, RTK ligands from the FGF, IGF, Neuregulin, and HGF families were predicted to signal to epithelial cells, and some were recently found to improve the phenotype of organoids grown from healthy intestinal epithelium^22,23^ or were linked to CRC EMT^24,25^. Other orphan interactions or cross-talks (e.g., TGFß/RTKs/Integrins/Selectins or Wnt/morphogens) also offer opportunities for further studies.

The many predicted missing cell-cell interactions led to some key scientific queries: Can the TME be rebuilt *in vitro* to recapitulate this complexity? Furthermore, would PDOs behave more similarly to OTs? Finally, are a few key factors responsible for most of the phenotypic drift, or is the TME essential in its entirety?

### Rebuilding the TME of CRC PDOs

To answer these questions, we worked on recapitulating the cellular TME with co-cultures of epithelial, stromal, and immune cells with a carefully selected matrix (**Figure S5A**). While immune cells are not reliant on the matrix, stromal cells thrive in collagen, but not Matrigel and PDOs need soft basement membrane-mimetic matrices. Previously, we had successfully grown intestinal organoids with PEG-RGD-laminin^26^, fibrin-laminin^27^, or on the surface of collagen: Matrigel 4:1^28^. Here, we verified that these conditions supported CAF cultures as well, with tuning of the collagen:Matrigel ratio for 3D encapsulation in the latter. We opted for collagen:Matrigel based on gene expression of PDOs (**Figure S5B-C**) and confirmed that immune cells could grow and migrate freely in this matrix (**Sup video**).

Regarding media, we observed that TIL cultures responded to activation by αCD3/28 mAbs in any base media with IL2 (**Figure S5D**). PDOs and CAFs had more stringent requirements, where higher serum content was supportive for CAFs but detrimental to PDOs, particularly under hypoxia (**Figure S5E**). To accommodate both, we opted for a 50:50 mixture of CAF and PDO base media, which had 2.5% serum. We also added GM-CSF to support in situ Mo-APC differentiation. We reasoned that further polarization of the APCs to various kinds of MoDCs or TAMs would be guided by other cells in the CRC+TME cultures. These cells were the only ones not proliferating in culture and hence overgrown by the rest of the cells in the absence of a constant supply from the bloodstream, which is a limitation of this study. Finally, as all cell types tolerated or benefited from hypoxia, we performed the PDO+TME cultures under physiologically relevant 5% O_2_.

### scRNA-seq profiling of co-culture arrays illuminates intercellular influence in CRC tumors

Eliminating a specific cell type to understand its impact is common in murine studies, but unfeasible in the context of human tumors. We reasoned that we could achieve the same purpose with selective reconstruction. We profiled the response of PDOs to TME cell types together and separately (**Figure 6A**). To this end, we used hashtag-based condition multiplexing, where such datasets are free of technical differences, to enable extremely robust differential expression analysis. To increase the profiling throughput, we combined this approach with genetic multiplexing of up to 4 patients, using a tailor-made gate-based demultiplexing workflow available in burgertools.

**Figure 6.**
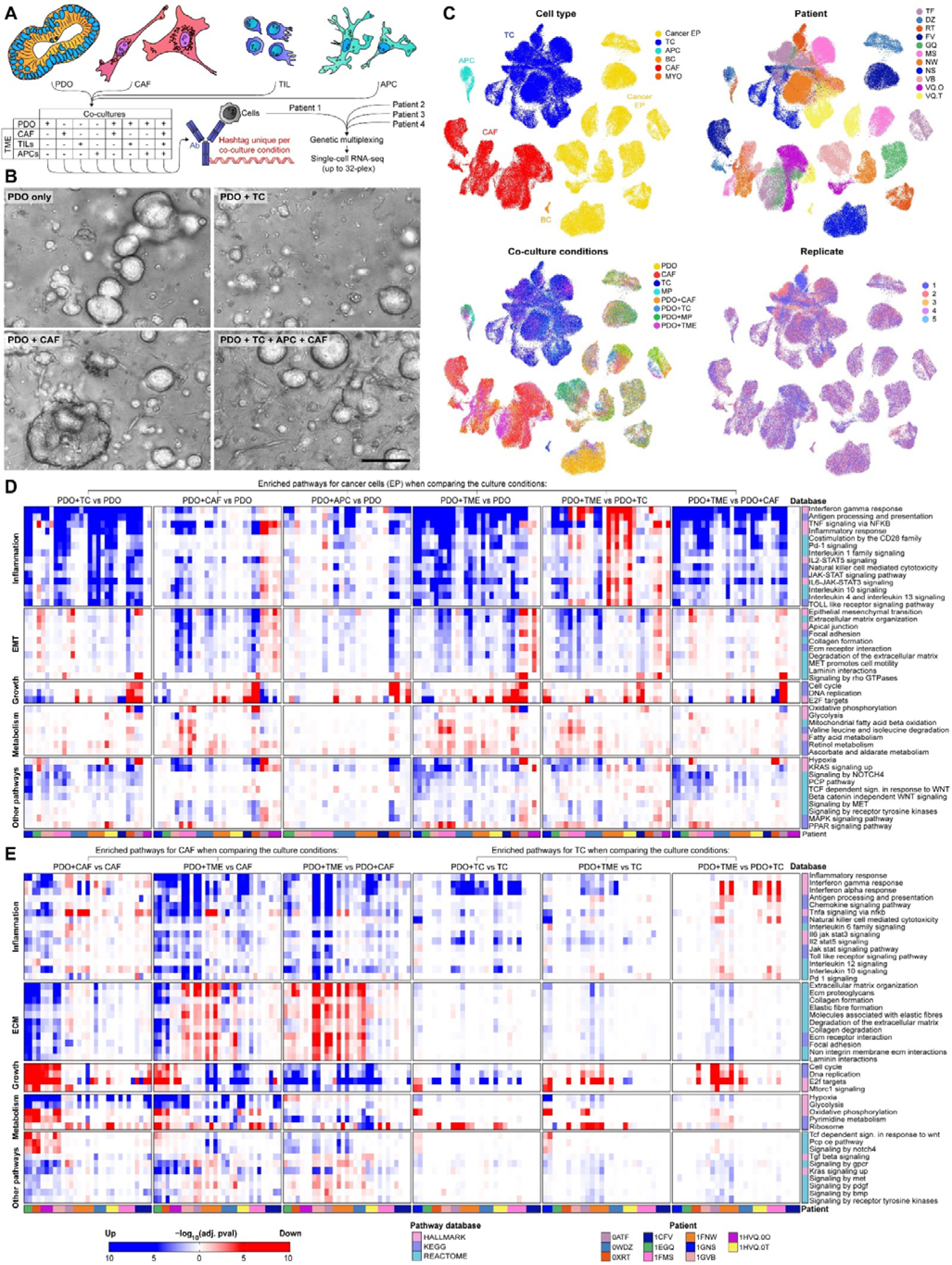
(A) Co-culture multiplexing schematic, with a combination of CITE-seq for co-culture conditions and genetic multiplexing of patients. (B) Examples of co-cultures in brightfield microscopy. Scale bar: 100 μm. (C) UMAP highlighting identities and replicates among the cells profiled as part of the co-culture datasets. (D-E) Select top pathways enriched in cancer cells and CAFs/TCs respectively, by fgsea, via comparison of a co-culture (e.g., PDO+TC) to a reference culture (e.g., PDO), organized by overarching category (left).

We performed time-lapse monitoring of the co-cultures and demonstrated that the cells could freely interact. The PDOs formed scattered blobs, and occasionally protrusions, while CAFs formed a coarse mesh with marked contractile activity that spread through the matrix (**Figure 6B**). Mo-APCs and TILs were highly motile and interacted with CAFs or PDOs (**Sup video**). In total, we acquired 260 independent datasets after demultiplexing, for a total of 182 k cells and 11 patients (**Figure 6C**). This *in vitro* PDO+TME data can be explored online.

We demonstrated through pathway enrichment analysis that cancer cells responded to various cell types in the TME alone or in combination (**Figure 6D**). The absence of significant pathway enrichment may sometimes be from insufficient data strength, and we therefore focused on the strongly significant or consistent enrichments. In summary, co-culture with TCs markedly increased diverse inflammation-related pathways, including IFN, TNF, and diverse interleukins (6D top left sub-panel). Some patients had a strong reduction of cell cycle-related pathways. Co-cultures with CAFs increased EMT-related transcriptional programs, with varied effects on proliferation. APCs were pro-inflammatory and pro-EMT, albeit to a lower extent than TCs or CAFs. Full co-culture yielded strong increases in both inflammatory and EMT signatures, with the greatest reduction in metabolism and growth-related pathways. Additionally, we noted the activation of various epithelium-controlling pathways such as KRAS, NOTCH, WNT, PCP, RTKs, and MET. Interestingly, a comparison of cancer cell gene expression with full TME vs. only TCs showed a marked reduction in inflammatory signatures, particularly interferons, in most patients, which suggests that the TME is immune suppressive.

With these same datasets, we could see how TME cells were affected by PDOs and the rest of the TME (**Figure 6E**). In the presence of PDOs, CAFs transitioned from a proliferative to a matrix-depositing phenotype. Numerous changes in pathway activations (e.g., TGFß, GPCR, KRAS, Met, PDGF, BMP, RTKs up, WNT/PCP, and NOTCH down) might explain this radical change. Interestingly, when we added additional immune cells, the CAFs showed an inflammatory response and switched again to a proliferative rather than matrix-depositing phenotype.

Among the immune cells, APCs were the only non-proliferative cells. Their low proportion at the end of the co-culture hindered significant enrichments and they were excluded from this analysis. Conversely, TCs responded to PDOs, marked by increased activation, inflammatory pathways, and proliferation. Consistent with the enriched pathways in PDOs, we found that including the rest of the TME reduced TC activation (6E top right sub-panel).

While this study focuses on key pathways conserved across patients, the entirety of the data is available in the online atlas.

### TME reconstruction rescues the gene expression drift between OTs and PDOs

An essential query was whether reconstruction of the TME could induce *in vivo*-like behavior in cancer cells. Toward this goal, we compared the gene expression changes induced by TME reconstruction to differences between in vivo and *in vitro* (**Figure 7A**).

**Figure 7.**
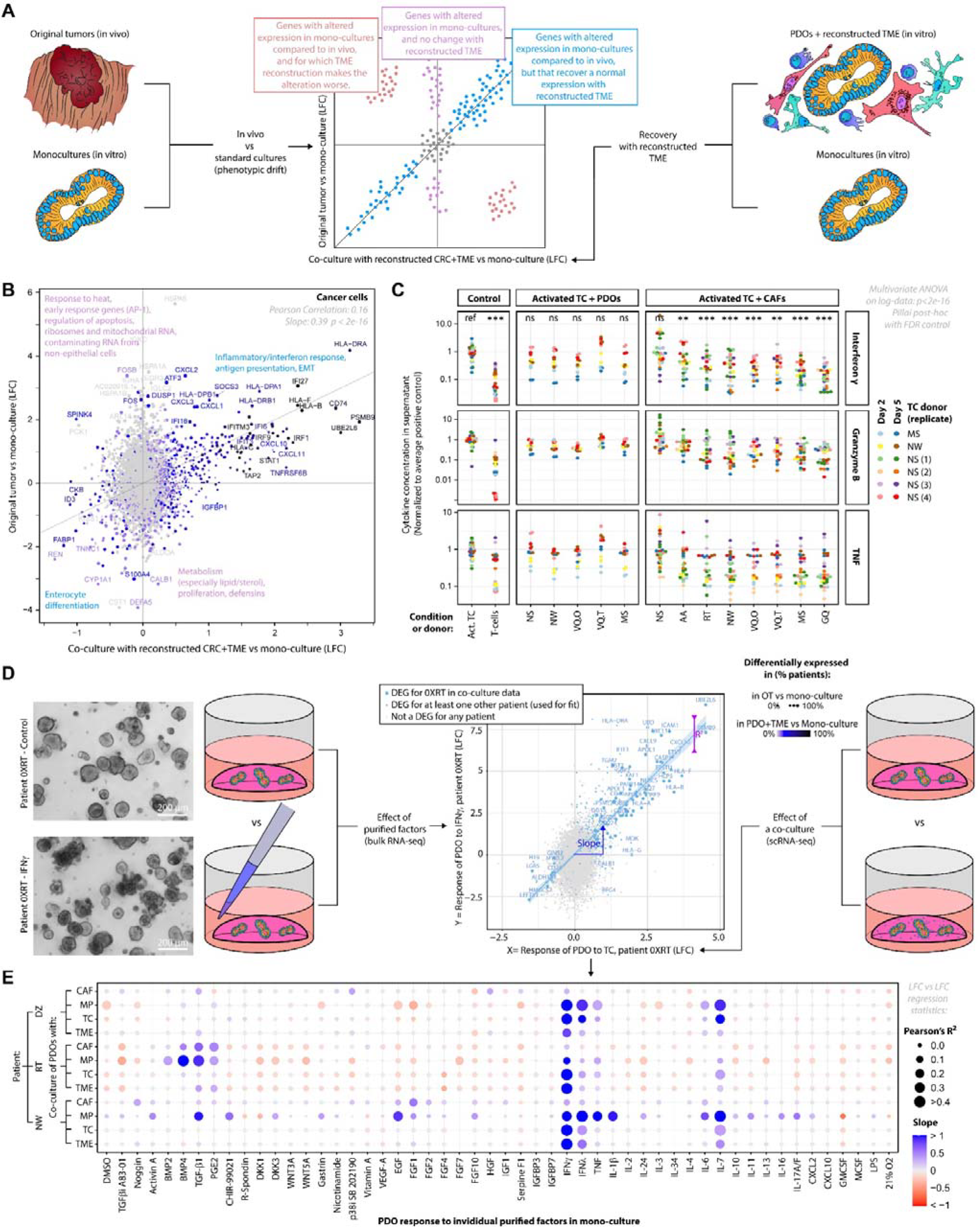
(A) Explanatory schematic of LFC-LFC plot. (B) LFC-LFC plots of cancer cell responses to in vivo TME vs. reconstructed TME. Average LFCs across replicates per patient, then across patients, for 16 patients in OT vs. PDO and 11 patients for reconstructed TME vs. monoculture. For co-cultures, genes significant in at least two scRNA-seq replicates for a given patient are defined as DEGs for the patient. Significance thresholds for individual comparisons, in co-cultures: p.adj<0.01 and FC>1.5; in OT vs. PDO (noisier): p.adj<0.01 and FC>2.38. (C) TC inhibition by CAFs and lack of inhibition by PDOs as demonstrated by functional assay with secreted cytokine CBA readout. Multivariate ANOVA post hoc compares each condition to the activated TC control. (D) Explanatory schematic of LFC-LFC plot linear fit, for correlation analysis of the response to a cell type vs. a purified factor. (E) Dot plot showing the slope and coefficient of determination that corresponds to each comparison of the cancer cell response to a reconstructed TME vs. a purified factor.

We found a strong correlation via log fold change (LFC) comparison plots (**Figure 7B**), which showed that the reconstructed TME recapitulated *in vivo*-like cell behavior. Strongly deviating genes mainly were related to early response, apoptosis, or mitochondrial RNA, all of which are associated with lower cell quality. It is anticipated that cells sequenced from an easily dissociated culture with immediate processing are less stressed than a surgical sample subjected to lengthy transfer and harsher processing. Therefore, we regarded these genes as technical artifacts rather than poorly recapitulated gene expression programs. Most other poorly recapitulated genes appeared to originate from environmental RNA contaminants from other cell types in the TME and were more present in tumor samples, another likely technical artifact. Thus, the quality of the phenotype recovery upon TME reconstruction is all the more striking.

DEFA5 and SPINK4 were notable exceptions, with substantial deviations and unlikely to be artifacts. They could reflect an abnormal tendency towards a Paneth-like polarity instead of Goblet-like polarity *in vitro*, which was not rescued in co-culture.

This data showed that cell proliferation and metabolism were at least partially recapitulated. Furthermore, the most striking changes, namely inflammation and EMT-related genes, were particularly well recapitulated. While the data shown here is averaged across patients to look for the most critical generic trends, full custom explorations are possible in the CRC-TME atlas.

### Co-cultures partially rescue TC but not CAF phenotype from in vivo

We repeated the analysis on TCs (**Figure S6**, left panel), which showed excellent recovery of genes related to TC activation and exhaustion/immune inhibition. However, outliers included markers of TC specialization to T_fh_, T_h_2, or innate-like lymphoid cells such as γδ/NKT, highlighting the loss of finer specialized populations.

On the contrary, the CAF phenotype (S6 right panel) was poorly recapitulated even with TME reconstruction, with no positive correlation. Strong outliers included relevant cytokines, growth factors, and matrix components. This highlights that CAFs are extraordinarily diverse and highly sensitive to their environment, which will need further study. However, the recovery of PDO phenotypes shows that cultured CAFs already reproduce the main effects they would have on cancer cells in vivo.

### CAFs are responsible for TC inactivation in CRC tumors

From the CRC-TME atlas, we made the striking prediction that CRC immune escape occurs through TC inhibition by other TME cells rather than direct TC inhibition by cancer cells. Since APCs primarily expressed pro-inflammatory cytokines, we hypothesized that CAFs were responsible for this effect.

We co-cultured TCs artificially activated with anti-CD3/CD28 antibodies with PDOs or CAFs, and simultaneously monitored inflammatory cytokines at the protein level (**Figure 7C**). We performed these assays in suspension cultures, which favored contact between TCs and other cells. We found that seven of the eight CAF lines significantly inhibited TC activity, but none of the 5 PDO lines tested. This confirmed that CAF rather than cancer cells are responsible for TC inhibition.

### Response profiles to individual factors suspected of mediating PDO/TME interactions

We functionally tested the most important candidates for cell-cell interactions by comparing the response of PDOs to co-cultures vs. over 50 individual purified factors, which were selected based on our predicted interactions, together with literature data and current organoid culture protocols (**Figure 7D, E**). The response to individual factors was profiled with a recently described highly multiplexed bulk RNA-seq method (BRB-seq^29^).

The response of PDOs to TCs was strongly correlated to interferon response, particularly IFNγ, as well as IFNλ and IL-7. The reaction to APCs correlated again with IFNs (albeit to a lesser extent), but also TNF, TGFß, IL-6, and IL-1ß. We observed correlations with BMPs and PGE2 in one patient and Activin A, IL-11, 16, 17A/F, CXCL2, and EGF in another.

Most interestingly, the CAF effect, attributed to TGFß in murine studies, is heavily correlated with FGF1 in the patient with the most robust response (and, to a lower extent, other FGFs and RTK ligands). This factor is not among the usual suspects in the CRC EMT literature, which makes it particularly interesting, and our findings are corroborated by another recent study^25. 25^IL6 and Noggin also showed weaker correlations. Another patient showed a more traditional correlation profile, with TGFß, BMP4, and PGE2 as best matches for the PDO response to CAFs. Finally, the response to full co-cultures was dominated by IFNγ, due to the high expression of interferon-stimulated genes, though other responses were present among lower expressed genes. While these factors are described in the literature, such direct comparison and simultaneous visualization of their strength in a well-defined system is unique and clarifies their relative importance in the field of systems biology.

## Conclusion

In this study, we demonstrated that reconstruction of the CRC-TME could improve the PDO phenotype, primarily by recapitulating the inflammation, and EMT-like behavior present *in vivo*. We identified the factors primarily responsible for these effects, namely IFNγ/λ with contributions from TNF, IL1ß, IL7, IL6, and FGFs, including potently FGF1, or combinations of TGFß with GFs rescuing proliferation. Proliferation is influenced by both in opposing directions, with overall slower growth *in vivo* and in PDOs with TME. Since many chemotherapies target proliferating cells, accurate reproduction of the right balance could greatly enhance the predictive power of PDOs. Importantly, in our work on TME reconstruction, we discovered that immune evasion was predominantly mediated by CAFs rather than cancer cells or APCs, and we suggest that CAF might have an equal leading role *in vivo*. Future work will be needed to clarify the mechanisms of this interaction, which is highly relevant to immunotherapies against CRC.

The extensive CRC and TME interaction data collected, made easily accessible with the CRC-TME atlas, is an expansive and informative knowledge base that can benefit future studies. We envision that our new bioinformatic pipelines, gathered in the package burgertools for reusability and the living biobank we established, can be highly applicable resources to facilitate such future endeavors.

The research presented herein could serve as a starting point for numerous new potential studies. While we did not recapitulate the vasculature, this is the next frontier in full TME reconstruction, and its inclusion would form highly cellular constructs without nutrient accessibility issues. Vasculature inclusion would facilitate the study of anti-angiogenic-drugs and recapitulate more biological interactions, including the pericyte niche and immune cell circulation. Furthermore, lymphocyte and fibroblast populations were tremendously more diverse and better differentiated *in vivo* than *in vitro*. We anticipate that understanding and developing control over these differentiation trajectories will soon be a major endeavor. Single-cell omics also provide a plethora of new biological predictions, from transcriptional networks to cell communication axes and unexpected pathway cross-talks. While we have functionally investigated the most prominent results, there is much more experimental groundwork to be covered by functional assays.

Conclusively, the rich and intricate data on the PDO/TME interactions provided herein will support research on new therapeutic avenues. Additionally, the refinements we introduced for the *in vitro* modeling of CRC will directly empower pre-clinical development and renewed efforts in personalized medicine.

## Acknowledgements

The project “Engineering and clinical validation of scalable organoid models for personalized oncology (OPERON)” was financed by the Personalized Health and Related Technologies initiative of the ETH domain, grant 2017-204, EPFL SCR0032439. We are thankful to the EPFL Gene Expression Core Facility (GECF), in particular Dr. Bastien Mangeat, the EPFL Bioimaging and Optics Platform (BIOP), and the EPFL Flow Cytometry Core Facility (FCCF) for their assistance with data acquisition.

