## Supplementary Data for "Dissection and reconstruction of the colorectal cancer tumor microenvironment"

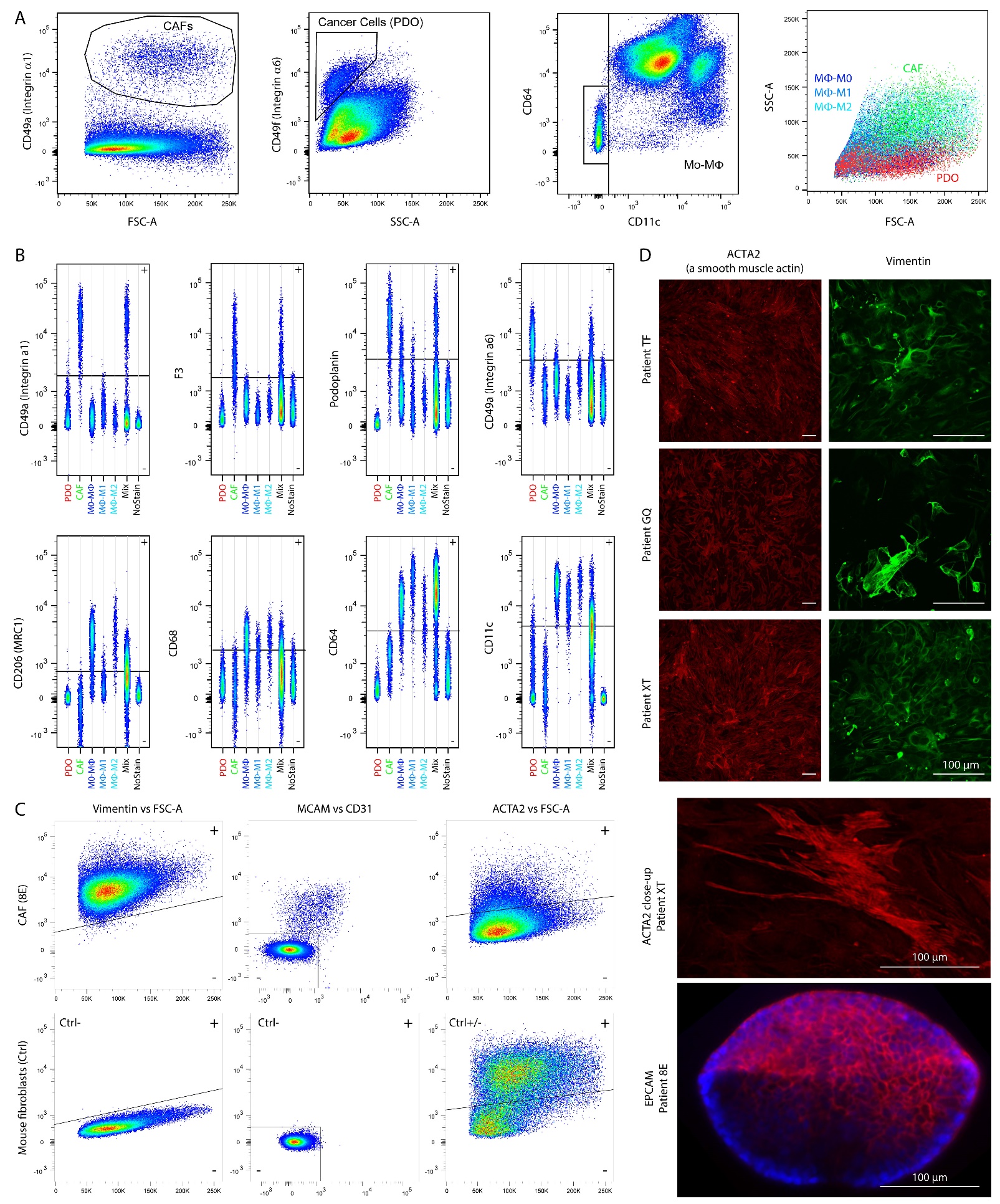

Figure S1. (A) Cell types in the TME can be distinguished by their surface markers in FACS scatter plots. (B) Surface markers associated with PDOs, CAFs and monocyte-derived macrophages (Mo-MΦ) cultures. M0-MF refers to monocytes only exposed to GM-CSF to acquire a generic unspecialized APC phenotype, M1-MF further are exposed to LPS, and M2-MF are exposed to IL4+IL10 instead. (C) Cells in the stromal cultures (CAFs) are positive for the intracellular marker VIM, and have varied amount of ACTA2, and have some residual MCAM^+^ CD31^-^ cells (undifferentiated pericytes). Mouse small intestinal mixed fibroblasts are used as a negative control for the human specific antibodies against VIM and MCAM, and positive control with positive and negative populations for the cross-species ACTA2 antibody. (D) Immunostainings of CAFs in 2D culture for ACTA2 and VIM and a retrieved PDO from 3D culture for EPCAM.

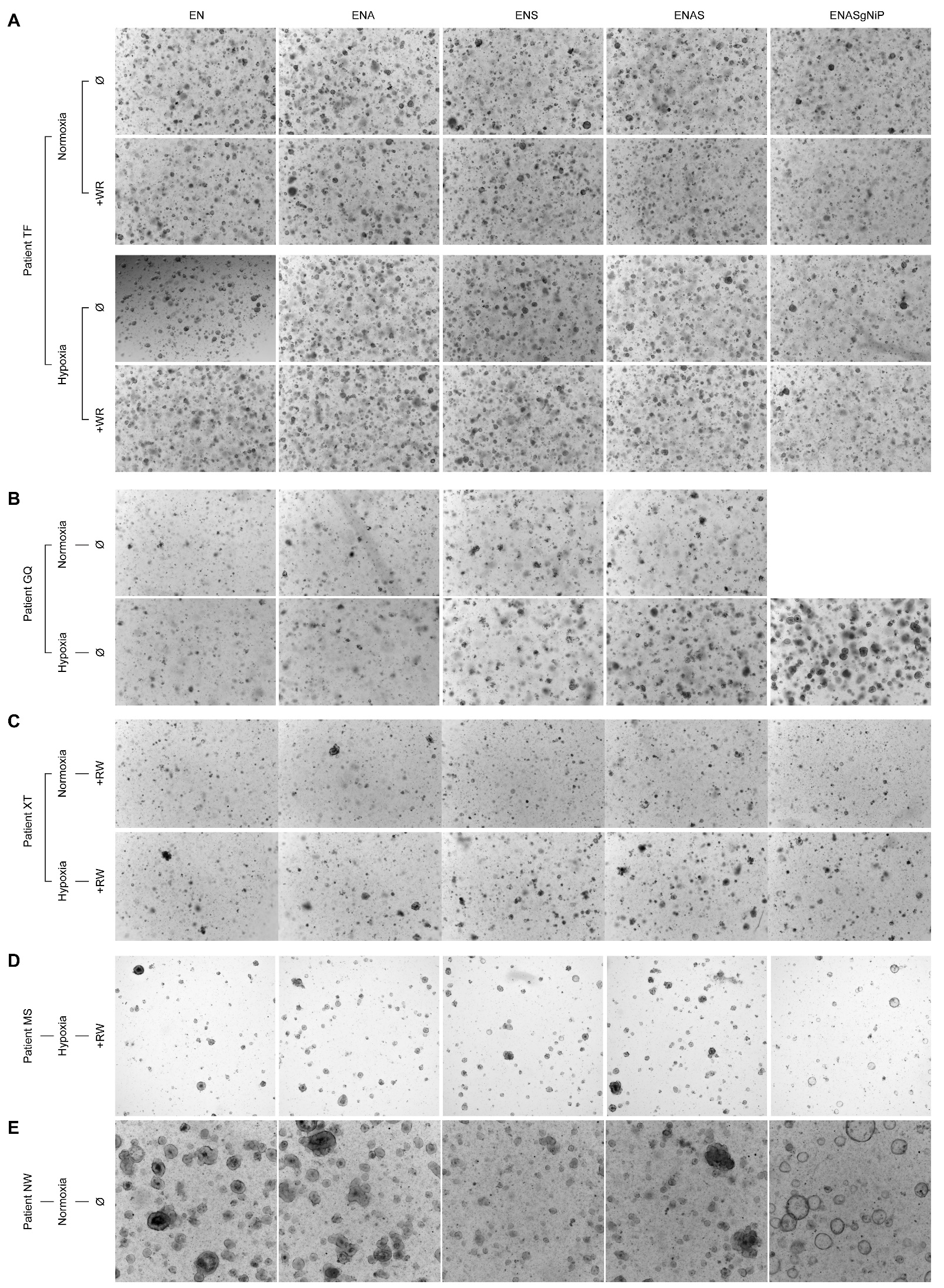

Figure S2. (A-E) Brightfield images of CRC PDOs during media screens at P0. (A) equal growth in all media and oxygen tensions. (B) Increased growth with richer medium. (C) Increased growth under hypoxia. (D) Reduced cell numbers and altered morphology with Prostaglandin-E (and nicotinamide). (E) Reduced growth in the presence of S (TGFß inhibitor).

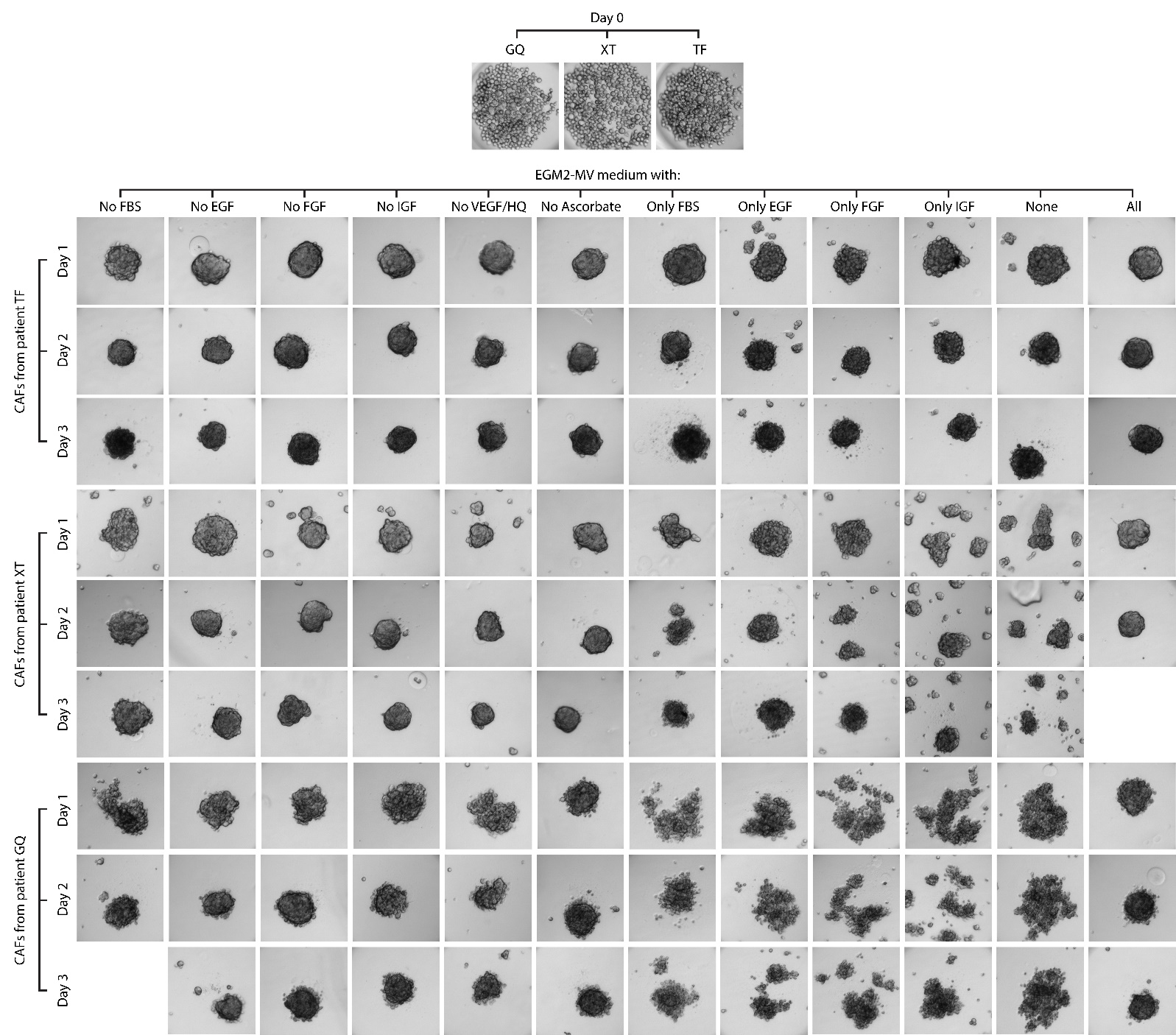

Figure S3. Brightfield images of CAFs in microwell suspension cultures (Gri3D^TM^, SUN bioscience), grown with EGM2-MV medium without individual factors, or with only individual factors, to determine which ones are essential. CAF health is judged by their potential to aggregate into a tight sphere and lack of dead cell / debris shedding, and is best when all factors are included for the three donors tested.

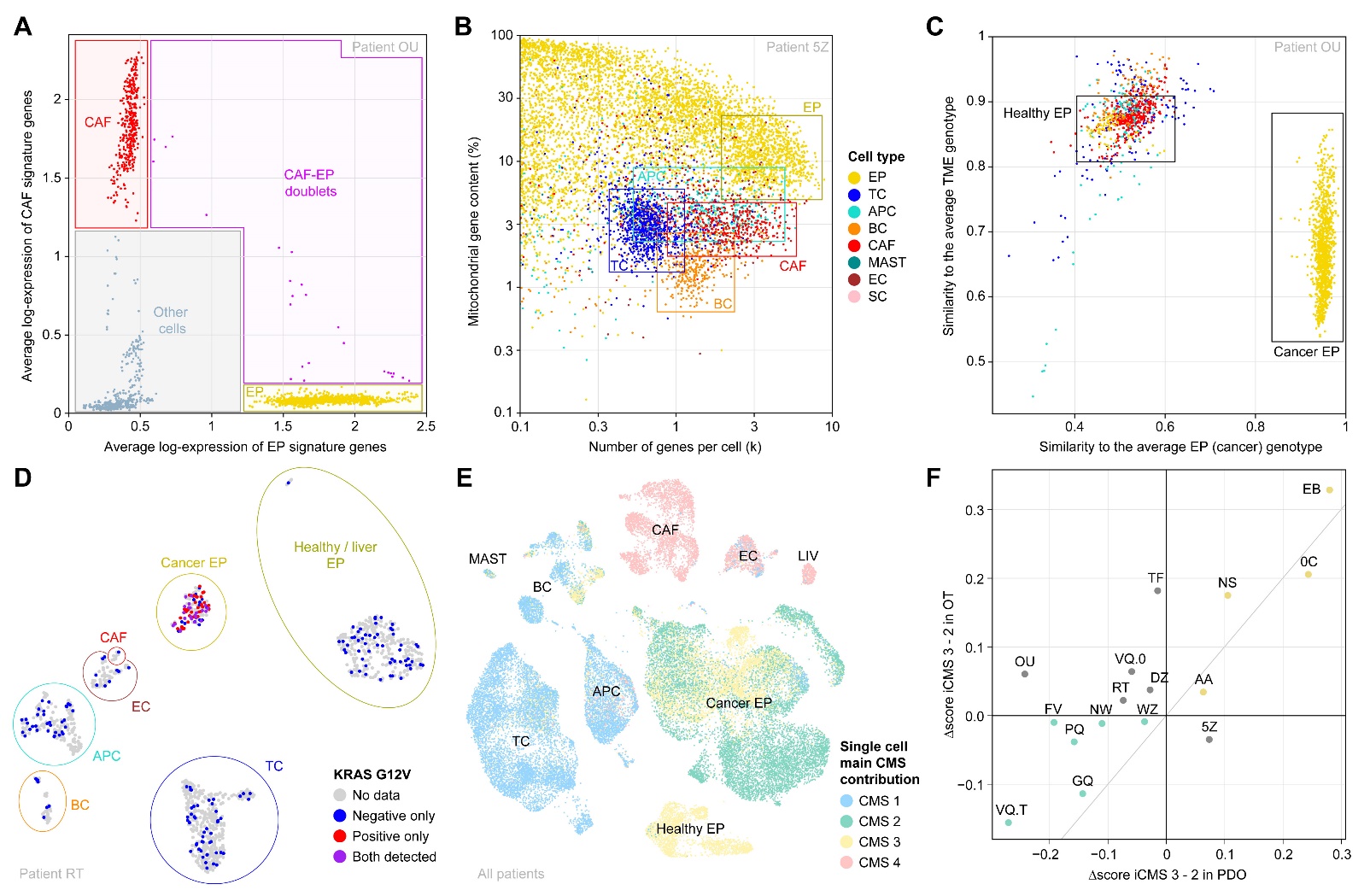

Figure S4. Single cell RNA-seq processing. (A) Cell classification by scoring cell type specific signatures at the single cell level and gating. In this illustration, a 2D scatter plot of single cells for the CAF and EP signatures is shown for simplicity, but gating is done simultaneously on all cell type signatures. Cells which pass several or no gates are tagged for manual examination, and upon confirmation excluded as multiplets/debris. Alternatively to manual gating, the cells can be assigned to the cell type for which the signature scores highest – a process we use before QC to get a first assignment, refined with manual gating afterwards. (B) Cell type aware QC. QC is done by gating on mitochondrial content and number of detected genes in single cells. The areas containing high quality cells of various types are highlighted. For example, live TCs tend to have the same gene number as dying cancer cells, whereas BCs and CAFs are similar to cancer debris in mitochondrial content. Doing the filtering in bulk as is customary would therefore leave a considerable amount of dead cells and debris, or exclude a considerable amount of TME cells, which would unnecessarily introduce a large bias in subsequent analyses. (C) Gating of cancer vs healthy epithelium. We scored the polymorphic sites within the patient pool, obtained from WES of PDOs, at the single cell level, and computed average genotypes of TME cells vs epithelial cells. We then restrict these genotypes to sites which are significantly different between the EP and TME, and compute the similarity (percentage matching) of single cells to these two reference genotypes. If the scores are too noisy for direct gating (e.g. too few variants significantly different between cancer and healthy EP, or too low sequencing depth), we used “cross-imputation”, defined as diffusion of the similarity scores between nearest neighbors in RNA space (cancer and healthy cells have very distinct gene expression). The resulting scores are used to construct scatter plots like the one displayed. Cancer cells cluster very distinctly from healthy epithelial cells, with higher similarity to the “cancer” (EP) average genotype and lower similarity to the TME average genotype, which enables easy gating of cancer vs healthy EP. The TME cells cluster together with the putative healthy epithelial cells, which gives a first validation of the approach. If cancer cells are outnumbered by healthy EP cells (which normally is not the case in tumor samples), the approach can be adapted by adding an additional step, separating subclusters of EP cells to define additional reference genotypes. (D) Examination of variant data at the single cell level can be used to further confirm the cancer vs healthy cell classification. For example, KRAS is a short gene which makes its mutations easily detected in scRNA-seq data. (E) Contribution of various cell types to the bulk CMS scores. Even though CMS classifiers are only meant to be applied on bulk RNA-seq, applying them at the single cell level shows how various cell types influence the overall score. (F) Correlation between iCMS (defined as CMS scored on pseudo-bulk RNA-seq cancer cells only) in vitro vs in vivo.

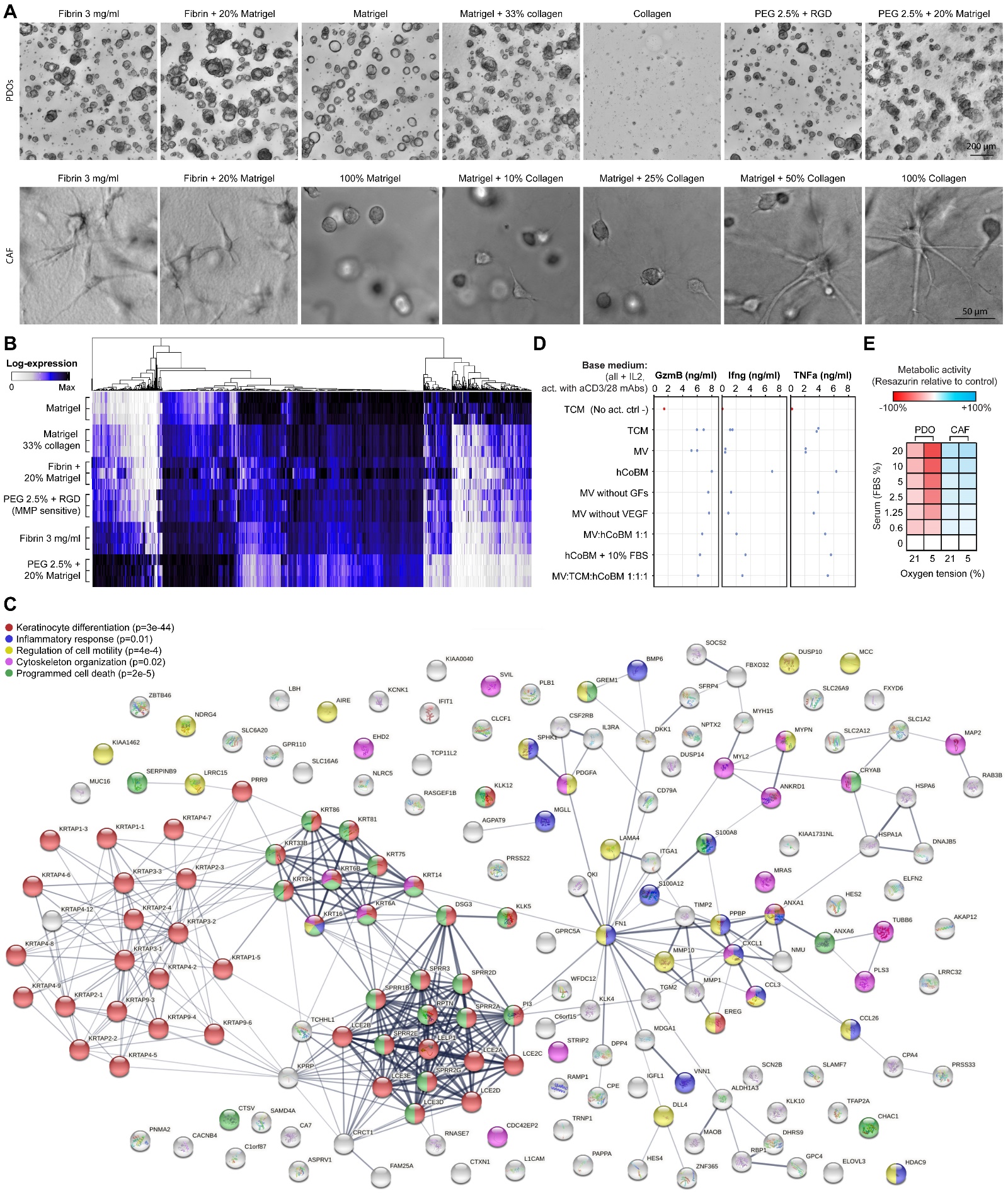

Figure S5. (A) Brightfield images of PDOs and CAFs cultured in different matrices. CAFs fail to spread in pure Matrigel, and PDOs die in pure collagen. Fibrin, PEG and Matrigel:Collagen hybrids support both cell types. (B) Heatmap of variable genes from RNA seq of PDOs grown in various matrices. Almost all genes have a continuous gradient from top to bottom. (C) Stringdb network and GO-annotations of genes upregulated in matrices other than Matrigel. The network is dominated by keratins typical of keratinocytes rather than intestine, not expressed CRC in vivo, highlighting that gene expression closer to that in Matrigel is more similar to original tumors. (D) Confirmation that T cells can be activated in various base media relevant to co-cultures. Cytokine concentrations in supernatant measured by CBA. TCM: T cell medium (RPMI+10% FBS based). hCoBM: human colon base medium (aDMEM/F12+B27 based). MV: Endothelial Growth Medium 2 – Microvascular. (E) Fetal bovine serum (FBS) inhibits promotes CAF growth. Average from n=3.

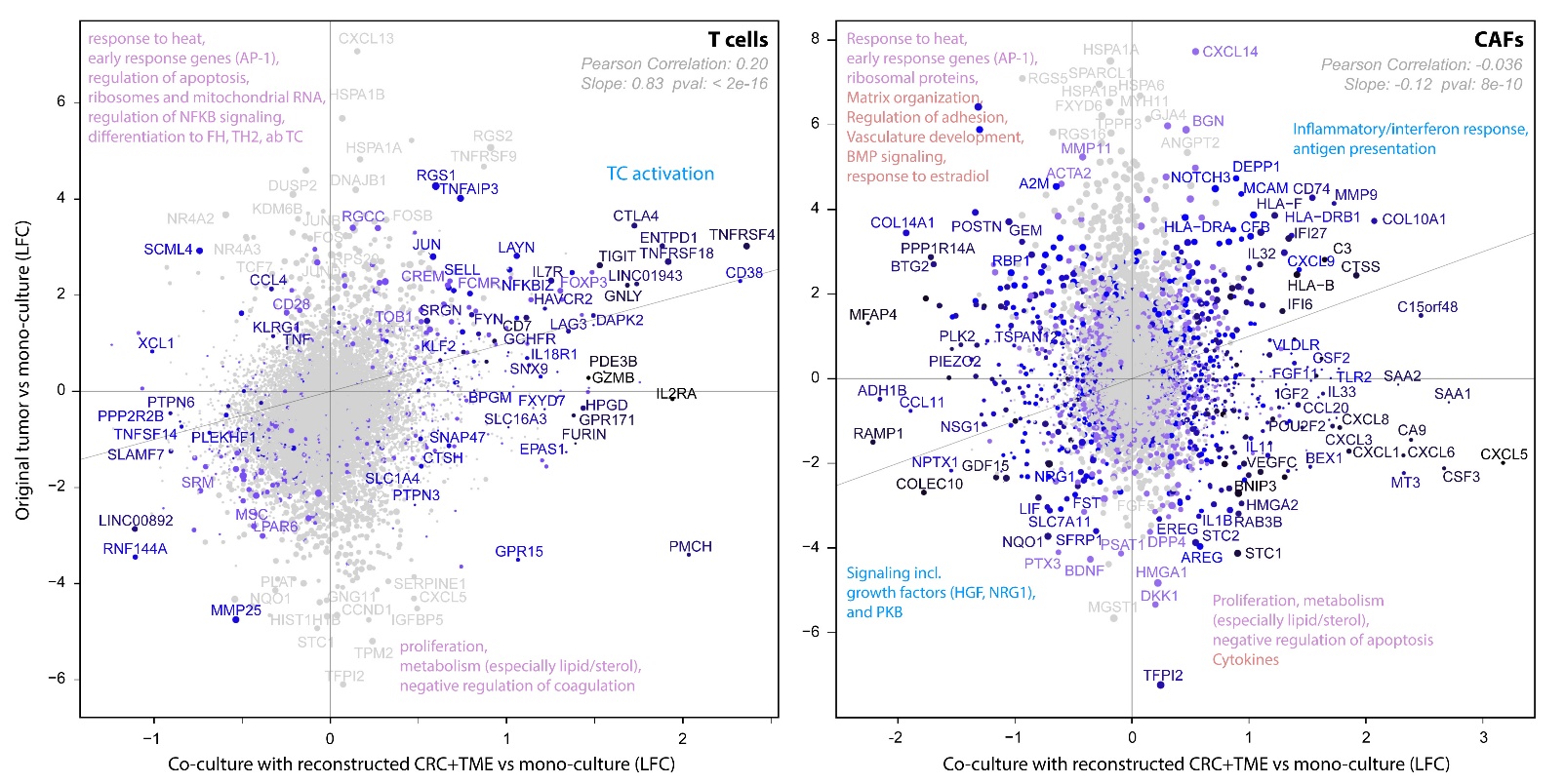

Figure S6. LFC-LFC plots of the response of TCs and CAFs to in vivo TME vs reconstructed TME. Average LFCs across replicates per patient, then across patients, for 9/10 patients in OT vs TC/CAF and 10/10 patients for reconstructed TME vs monoculture respectively. For in vitro TME reconstruction, genes significant in at least 2 scRNA-seq replicates for a given patient are defined as DEGs for the patient. Significance thresholds for individual comparisons, in co-cultures: p.adj<0.01 and FC>1.5; in OT vs PDO (noisier): p.adj<0.01 and FC>2.38.

**Materials and methods**

*Sample collection*

Patient samples were collected by the Centre Hospitalier Universtaire de Lausanne (CHUV) with informed consent under the protocol CHUV_DO_CTE_TRP_0001_2017 - Tumor MicroEnvironment characterization and ex-vivo immune intervention.

*Tumor dissociation*

Tumor surgery samples or needle-core biopsies were collected in saline and transported on ice. They were then cut in small pieces with a scalpel and rinsed three times with ice cold basal medium (BM), which consist of advanced DMEM-F12 (Gibco 12634-010) + 1x Glutamax (Gibco 35050-087) + 1x HEPES (Gibco 15630-122) + 1x Pen/Strep (Gibco 15140-122), followed by digestion in 10 ml of collagenase I (Gibco 17100-17) at 1.5 mg/ml in BM for 30 min and 0.1 mg/ml hyaluronidase (Sigma H3506) at 37°C with shaking. After trituration with a 10 ml serological pipette, the remaining fragments were collected by filtration on a 100 µm strainer, and further digested with TrypLE (Thermo 12605028) for 10 min at 37°C with shaking, triturated with a 10 ml Pasteur pipette and strained again. The combined lysates were washed 3 times with 10 ml BM+10% fetal bovine serum (FBS, Gibco 10500-064), and once in BM, using centrifugation at 300 g for 4 min for collection, then resuspended at approximately 1 M/ml in BM. Any remaining tissue pieces when applicable were further digested for 2h with collagenase II (Gibco 171-01-15) 1.5 mg/ml in BM at 37°C with shaking, and the resulting cells and fragments were washed similarly but not merged with the rest, and exclusively used for stromal/CAF cultures. The typical yield was of 100-500 k cells from a needle core biopsy, and 5-30 M cells from a surgery sample.

*Monocyte isolation*

Samples of 7 ml of full blood were diluted 2-fold with PBS (Gibco 10010-015), and 7 ml were layered on top of 7 ml of lymphoprep (Stemcelltech 07851) in two falcon tubes. They were then centrifuged at 1000 g for 30 min with brakes off. The peripheral blood mononuclear cells (PBMCs) were collected at the interface between the upper/yellow and middle/clear phases and transferred to a new 50 ml falcon tube, in which they were washed with ~45 ml of PBS+10% FBS and collected by centrifugation at 500 g for 10 min. The cells were picked up in 100 µl of PBS+0.5% BSA (Sigma 10735086001), and the CD14^+^ monocytes were isolated by magnetic activated cell sorting (MACS) according to the manufacturer protocol (Miltenyi 130-050-201). Typical yield was of 4-10 M cells.

*Growth factors for patient derived organoid (PDO) culture*

The factors used in PDO culture (and their denomination in abbreviated media composition designations, concentration in medium, and origin) were: EGF (E, 50 ng/ml, R&D 236-EG-200), Noggin (N, 100 ng/ml, EPFL protein production facility), A83-01 (A, 500 nM, Stemgen/Reprocell 41030), SB202190 (S, 10 nM, Selleckchem S1077), gastrin (g, 10 nM, Sigma G9145), nicotinamide (Ni, 10 mM) Calbiochem 481907), prostaglandin-E2 (P, 10 nM, Sigma P6532), R-Spondin (R, 500 ng/ml, EPFL protein production facility), and Wnt3a (W, 100 ng/ml, Time bioscience rmW3aL-010).

*PDO culture*

Cells from tumor digest were resuspended in Matrigel (Corning 356231) at approximately 1 M cells/ml, and plated in a pre-warmed 24 well plate at 37°C in 20 µl domes in the middle of the wells. The plate was then flipped upside-down and returned to the incubator for 15 min gelation. As a standard, 12 domes per plate in two plates were done, with each well exposed to a unique culture condition for initial screening of media. The human colon base medium (hCoBM) consisted of BM + 1x B27 without vitamin A (Gibco 12587-001) + 1 mM N-acetylcysteine (Sigma A9165) + Primocin 1:500 (50mg/ml stock, invivogen ant-pm-02). Each plate had the media hCoBM, and hCoBM + EN, ENA, ENS, ENAS, ENASgNiP, both with and without RW, and one plate was kept in a normoxia incubator (21% O_2_, 5% CO_2_, 37°C, humidified), the other in a hypoxia incubator (2% or 5% O_2_ in early trials, 5% O_2_ exclusively after early sequencing experiments showed it to be the most physiological). When the cells available were enough, the initial trials could be done in duplicate and additional conditions could be tested, whereas when the cells were insufficient (some needle-core biopsies), the conditions with RW and with no growth factors were omitted from the initial screen. If still insufficient, the cells concentrations were reduced accordingly.

Medium was renewed every 3 days or when changing color, and the results of the initial medium screen were imaged immediately before first passage, which was when some wells approached overgrowth, or media changed color in less than two days, or when Matrigel drops start to lose mechanical integrity because of extended time in culture with poor growth.

The PDOs were then kept in the minimal medium supporting robust growth, which most commonly was hCoBM or hCoBM+EN, but occasionally could be other media (**Figure S2**). In general RW produced no visible difference, highlighting that colon cancers acquire mutations such as APCmut that make them function independently of Wnt early on, as expected. Hypoxia was in general either better or equal to normoxia, and when both functioned, hypoxia was kept, as the environment is then more physiological.

For passaging, Matrigel domes were mechanically broken by trituration in their medium using a 1 ml pipette, followed by collection by centrifugation at 250 g for 3 min. After removing the supernatant and Matrigel+debris/dead cells layer, the PDOs were either treated with trypLE for 5-10 min (visual monitoring) for single dissociation, or treated with trypLE for 1 min and lightly triturated with visual monitoring for passaging as fragments. Fragments were preferred for cell expension, as PDOs regrow faster, whereas single cells were preferred for assays that require precise starting cell numbers (10 k cells per 20 µl gel) and quantification of the response to growth factors. When passaging as single cells, the initial medium is supplemented with 1 µM of the ROCK inhibitor thiazovivin (Stemgen/Reprocell AMS.04-0017).

*TIL culture*

Cells from the tumor digest were resuspended in T cell medium (TCM) consisting of RPMI 1640 with glutamax (Gibco 61870-010) + 1x Pen/strep + 1x HEPES + 50 nM beta-mercaptoethanol (Gibco 31350-010) + 10% FBS + 500 U/ml IL2 (Peprotech 200-02). For T cell activation, the IL2 concentration was increased to 5000 U/ml and the medium supplemented with 1 µg/ml of anti-CD3 (Biolegend 317326) and anti-CD28 (Biolegend 302934) monoclonal antibodies. T cells were activated immediately after being put in culture or thawed from stocks to promote recovery and fast expension, and otherwise kept in the resting medium (unless otherwise indicated in figure legends for the needs of particular inhibition experiments). The cells were seeded at approximately 10 k per well in round-bottom anti-adhesion 96 well plates with 200 µl of medium for routine propagation, and the medium was changed every 3 days. Passaging was done by collecting, diluting, and reseeding.

*CAF culture*

Cells from combined first/second and third tumor digests were resuspended in EGM2-MV medium (Lonza CC4147) and seeded in collagen-coated T25/T75 flasks. Collagen coating was performed by swiftly diluting collagen (Atelocell IAC-50, native collagen from bovine dermis in solution at 5 mg/ml pH3.0) 500 fold in ice-cold PBS, then adding enough to completely cover the flask bottom and incubating at 37°C for at least 10 min and until usage. Before seeding, the PBS was aspirated and the cells in medium added immediately without letting the coating dry. The cells were passaged 1:3 when reaching confluence, by washing with PBS, detaching with trypsin-EDTA (Gibco 25200-072) while monitoring and tapping the flask every 2 min, blocking with BM+10%FBS, and resuspending in EGM2-MV for reseeding. For highly confluent flasks or for FACS analysis, trypsin was occasionally insufficient for complete dissociation because of the large amount of collagen fibers deposited by the cells, which was resolved by additional treatment of the collected cell sheet with collagenase I at 1.5 mg/ml in BM for 10 min before washing and resuspending.

*Biobank storage*

For cryopreservation, PDO fragments or TIL/CAF/Monocyte dissociated cells were resuspended in recovery cell freezing medium (RCFM, Gibco 12648-010) and frozen to -80°C in a Corning CoolCell LX Cell Freezing Container, then transferred to liquid nitrogen storage.

*scRNA-seq profiling of original tumors (OTs) and mono-cultures*

A fraction of the cell suspension obtained from tumor dissociation was immediately processed for scRNA-seq. PDOs and CAF were dissociated to single cells as described for passaging for their profiling in the same manner. If needed, additional clean-up was performed: in rare cases red blood cells were numerous in OTs, they were removed by lysis (Biolegend 420301), or if cell multiplets were still present after enzymatic dissociation they were removed by straining at increasingly small strainer pore size (70, 50, 30 µm), with visual examination. The cells were then immediately profiled with 10X Genomics gene expression kits according to manufacturer instructions, targeting 1500-3000 cells per sample. The samples were processed with the 5’v1.1 version of the kits, with the exception of the few earliest (3’v2) and latest (5’v2) samples.

The resulting libraries were sequenced on Illumina by the EPFL gene expression core facility with a read length of at least 90/26, with approximately 30 k usable unique reads per cell after filtering, and aligned to the genome GRCh38 with cellranger 6.1.2.

*Testing of alternative ECMs*

For encapsulation in fibrin 3 mg/ml, single cells at 10x the desired concentration, fibrinogen at 30 mg/ml (Baxter Tissucol/Tisseel diluted in Hank’s balanced salt solution or HBSS, stored aliquoted at -80°C and thawed at 37°C), thrombin at 5 U/ml (Baxter Tissucol diluted in HBSS), calcium 100 mM, and hCoBM were combined in proportions 1:1:1:1:5, at room temperature, with fibrinogen added last. After fast mixing, 20 µl drops were cast in 1 mm thick PDMS rings adhered to the bottom of 24 well plates, and left to gel upside-down in a 37°C incubator for 5 min before addition of medium. For fibrin+20% Matrigel, the proportion of hCoBM was reduced accordingly, and the mixture was kept on ice until addition of fibrinogen.

For encapsulation in collagen, 400 µl of stock solution (Atelocell IAC-50, native collagen from bovine dermis in solution at 5 mg/ml pH3.0) were cooled on ice, supplemented with 10% 10xMEM (Gibco 12800-058) and neutralized by addition of sodium bicarbonate (Alfa Aesar/Thermo 144-55-8) while monitoring the phenol red color reporter (orange for pH 7.4), with rapid mixing while trying to avoid bubble formation. The cells were then resuspended in this solution and plated as for Matrigel.

For encapsulation in collagen:Matrigel mixtures, neutralized collagen in medium was prepared as above and mixed with ice-cold Matrigel, with the two solutions mixed in the proportions indicated in the figures (e.g. 10%, 25%, 33%, 50% of the collagen solution). The cells were then resuspended and plated as for Matrigel.

For encapsulation in PEG gels, we proceeded as described previously^1^, with 2 %(w/v) 4-arm-PEG-VS content, cross-linked with MMP cleavable peptides Ac-GCRE-GPQGIWGQ-ERCG-NH2, with 1 mM of Ac-GRCGRGDSPG-NH2 as an RGD adhesion cue, in HEPES buffer pH8.0. For PEG+20% Matrigel, the volume of buffer was reduced accordingly and replaced with Matrigel, keeping the mixture on ice and adding the vinyl sulfonated PEG last. Of note, there is some presumably collagen IV aggregation in PEG-Matrigel (but not PEG-laminin) mixtures, which results in some fibrous inclusions in the gels, not seen in the other gel mixtures tested. The gels were cast in PDMS rings as for fibrin and incubated for 15 min upside-down before addition of medium.

The PDOs cultured in the various matrices for 10 days were released from the matrix with the adequate enzymes (Nattokinase from JBSL-USA for fibrin, TrypLE for Matrigel, Collagenase I for Collagen:Matrigel, Dispase from Thermo for PEG), collected in Trizol for RNA-isolation, and the transcripts sequenced with QuantSeq 3’ mRNA-seq FWD kits according to the manufacturer’s protocol, and sequenced on Illumina NextSeq by the EPFL gene expression core facility, with a target of 14.3 M reads per sample before filtering.

For analysis, the counts were normalized with log1p of counts per million. Differential expression and heatmap plotting of differentially expressed genes (DEGs) were performed in EdgeR 3.40.2 with a genewise negative binomial generalized linear model, using a threshold of 2 in log fold change and 0.001 in adjusted p-value for significance. We noticed all the DEGs were following the same trend with a difference of magnitude between matrices, rather than showing diverse signatures depending on the matrix, and used stringdb^2^ to annotate and interpret this gene signature.

*Co-culture*

Co-cultures were done in a Collagen:Matrigel 1:3 mixture. The cells were mixed with a target of 10 k CAFs, 20 k T cells and monocytes, and PDOs split as fragments ~1:3 to 1:5 depending on their density, for a full co-culture. For partial co-cultures, the cell numbers were the same but only relevant cell types were included. The mix was collected by centrifugation and resuspended in the ice-cold gel precursor mix, then plated as 20 µl domes in pre-warmed 24 well plates and returned to the incubator for 15 min for gelation. We then added co-culture medium, consisting of hCoBM:MV 1:1 supplemented with 3000 U/ml IL2 and 25 ng/ml GMCSF (R&D 215-GM-010). For PDO lines that need additional factors for survival, the factors were added during the initial culture to sustain growth, but removed in the last media change before sequencing in order to make cell-cell signaling apparent. The cells were then kept in culture until PDOs would have needed passaging, at which point they were analyzed. Time lapse-monitoring of the co-culture was performed in brightfield on a Nikon Ti microscope.

*scRNA-seq of co-cultures*

Co-cultures were performed with 3 to 6 individual gels (20 µl domes in 24 well plates) per dataset, and at least two replications of each dataset. Each dataset included 8 multiplexed conditions: all cell types grown in monoculture as controls, PDOs co-cultured with each TME cell type in isolation, and PDOs co-cultured with the full TME. The collagen in the co-culture matrix was first dissolved by replacing the medium in the culture wells with 0.5 ml of collagenase II 1.5 mg/ml in hCoBM for 30 min. The gels and supernatant were then collected with a 1 ml pipette, and centrifuged at 500 g for 6 min. The pelleted gel fragments were resuspended in TrypLE and incubated for 10 min to remove the Matrigel and dissociate the cells. We then blocked the dissociation by addition of 4 ml of BM + 10%FBS, collected the cells by centrifugation, and introduced the condition-specific hashtags (Biolegend Totalseq C 3-10 hashtagged antibodies), diluted 1:500 in FACS buffer. We washed twice with 5 ml of BM + 10% FBS, resuspended in 1 ml of BM + 10% FBS, merged the conditions and strained at 40 µm, collected by centrifugation, resuspended in a small volume of BM, and counted. As this step, the multiplexed cells from up to 4 patients were further merged for genetic multiplexing. The cell suspension was then processed with a 10X Genomics 5’ gene expression kit with feature barcoding (v1.1 for early samples, v2 for later samples).

Sequencing and alignment were then done as described above for OTs and monocultures.

*FACS*

For FACS analysis, monocytes/APCs were collected by incubation in ice cold PBS+EDTA and scraping, CAFs/stroma were collected by trypsinization + treatment by collagenase I as described above, and PDOs were dissociated with TrypLE as described above. Fc-receptors were blocked (Miltenyi 130-091-935). The cells were stained with CD49a, CD49f, CD64, CD11c, F3, Podoplanin, CD206, and CD68 for 30 min on ice. All antibody dilutions and washes were in FACS buffer consisting of PBS + 2%FBS + 5 mM EDTA (Invitrogen 15575020). For analysis of intracellular markers Vimentin and ACTA2 (alpha smooth muscle actin), the cells were fixed with paraformaldehyde 4% in PBS (Thermo 15434389) and permeabilized with saponin 0.1% (Sigma 47036). The cells were rinsed 3 times, and dead cells stained with AQUA (Invitrogen L34966), before resuspending in FACS buffer and straining at 50 µm. FACS experiments included cells killed by addition of methanol as a dead control, as well as unstained and live/dead only stained live controls, and either FMOs or control cells of other types to ensure antibody specificity and adjust gate levels. Cells were gated first for dead cell and doublet exclusion, then analyzed as shown in **Figure S1**.

*Immunocytochemistry*

Samples were fixed for 1h with 4% paraformaldehyde in PBS, blocked with PBS + 5% BSA overnight, incubated with primary antibodies in PBS + 5%BSA overnight, washed three times for >1h with PBS + 5% BSA, incubated with secondary antibodies in PBS + 5% BSA overnight, washed three times with PBS + 5% BSA, incubated for 1h with DAPI (Tocris 28718-90-3), then washed three with PBS, and imaged on a Leica SP8 confocal microscope. Antibodies used were EPCAM, ACTA2 and Vimentin.

*Mutation profiling by whole exome sequencing (WES) of PDOs*

WES was acquired by the Beijing Genomics Institute (BGI) by submitting to the DNBseq service with a coverage of >100x.

For mutation analysis, the reads (fastq files) from single WES datasets were aligned to the genome GRCh38 with bwa-mem 0.7.17^3^, and filtered and called with bcftools/samtools 1.14^4^ . The sam were sorted by read name with samtools sort -n, tagged for mate scores with samtools fixmate -m, sorted by coordinate with samtools sort, in case of several source fastq merged with samtools merge, deduplicated with samtools markdup -r, filtered for high quality and exported as a bam file with samtools -b -q 20, and indexed with samtools faidx. The variants in individual files were then called with samtools mpileup and bcftools call, high quality variants were filtered with vcftools --minQ 30, ID-annotated with bcftools annotate, and functionally annotated with the Ensembl variant effect predictor (VEP)^5^. The mutations highlighted in **Figure 1** are only those predicted to be pathogenic by VEP, on genes with a mutation prevalence of at least 1% in CRC according to the cancer genome atlas (TCGA, https://www.cancer.gov/tcga).

*Variant exploration at the single cell level*

For variant analysis in single cell data (used for demultiplexing, examination of cancer mutations apparent in single cells, and distinguishing cancer cells from neighboring healthy epithelium), filtered bam files from the previous WES analysis were reanalyzed together to highlight differences between patients and conserve rare variants. Genotype likelihoods were estimated, and variants jointly called on all datasets, with bcftools mpileup piped to bcftools call -m. We filtered high quality variants with vcftools -gzvcf --minQ 30 --recode --recode-INFO-all ^6^, and then compressed and indexed the joint vcf file with bgzip and tabix (repeated on all vcf files in following step). The variants were annotated with bcftools annotate, the chromosome naming conventions were changed for consistency with cellranger conventions with bcftools –rename-chrs, and multi-alleles were decomposed into a sum of monoallelic differences for compatibility with vartrix conventions while simultaneously fixing sites where the reference allele is not defined as the reference genome, with bcftools norm –multiallelics -any --check-ref. Finally, the variants were annotated and output in a tsv file with vep --tab --everything --nearest “symbol”.

Variants in scRNA-seq data were evaluated with vartrix v1.1.22 ([github.com/10XGenomics/vartrix](https://github.com/10XGenomics/vartrix)) based on the reference genome, list of cell barcodes, bam files output by cellranger, and final vcf file from the joint calling. The burgertools package provides tools to import of annotations in this format in R and to a Seurat object, and the interactive exploration of the variant information at the single cell level (c.f. following section*)*.

*scRNA-seq pre-processing*

The count matrices from cellranger were picked up in R and loaded in a Seurat v4 object. When variant data from vartrix or hashtags were available, they were loaded as additional modalities/assays. The RNA data was log-normalized with a sum of 10 k and a pseudo-count of 1, using natural logarithm.

We first performed an automatic pre-classification of the cells for the needs of quality control (QC) by cell type. The expression of most individual genes has a strong stochasticity and can be plagued by dropouts, so classifying single cells based on the expression of a few marker genes is normally not feasible. This is the rationale for classically going through a clustering step before defining cell types. We found that imputation is an excellent alternative, that enables us to reliably classify single cells based on the expression of even relatively simple gene signatures, without the need to define arbitrary clusters. We further found that MAGIC^7^ was excellent both performance-wise (which is essential to impute before QC) and in the quality and tunability of the filtering, even though we had to reduce the neighborhood of the diffusion compared to defaults in order to avoid carry-overs of gene expression between distinct cell types in small populations (knn=3, t=2, set as default in the burgertools MAGIC imputation wrapper). Key advantages of this approach are that classification can be done in a similar way to FACS, by gating, which can be monitored and adjusted. The delimitation between two neighboring populations in particular becomes a rational human decision, based on precise marker or signature ratios, rather than a stochastic division. Rare cell types, which are grouped with other cells by clustering algorithms and therefore missed in traditional workflows, are easily picked up with this approach, as they stand out clearly in signature scatter plots (we routinely detected as few as 1 to 3 cells of rare types such as Schwann or MAST cells in OT datasets). And automatic assignment of cells to the signature with the maximum score could easily be done in order to preview cell types before QC, even though clustering would be very impractical at this point.

We defined signature scores as the average of the log-transformed MAGIC-imputed RNA expression values of the genes in the signature. The signatures used for the first classification of cells into categories shown in Figure 1 and used for QC by cell category were:

T cell-like, including NK and other T-like native lymphocytes (TC): CD3D, CD3E, CD3G, CD4, CD8A, CD8B, GZMA, IFNG, NCAM1, KLRK1, JAK3

B cell-like, including plasma cells: IGHM, CD79A, CD79B, MZB1, DERL3, IGLL5, MS4A1, IGHD, IGHG1, IGHG3

APC-like, including monocytes, macrophages and dendritic cells: CD68, ITGAX, CD14, FCGR1A, MRC1, LYZ, FCER1G, TYROBP, S100A8, MMP9, C1QA, C1QB, C1QC, IGSF6, SPI1

Epithelial, including cancer and healthy epithelium: EPCAM, PHGR1, TFF3, CKB, AGR2, PLCG2, CDX2, PIGR, PHGR1, CD24, LGALS4, GPX2, CEACAM6, CLDN3, KRT8, KRT19, KRT18, TSPAN8, OLFM4, FABP1, REG1A

CAF-like, including fibroblasts and pericytes: LOXL2, TAGLN, COL6A1, ITGA1, MMP1, MMP3, ACTA2, COL1A1, CALD1, COL3A1, COL1A2, COL6A3, COL6A2, THY1, DCN, COL5A2, COL5A1, PDGFRB

Endothelial cells, including vascular and lymphatic: CD34, PECAM1, LYVE1, PLVAP, RAMP2, VWF, ESM1, FLT1, PODXL, GNG11, ENG, SLC9A3R2, HSPG2

Liver, including hepatocytes and ductal cells: ALB, CYP3A4, CYP3A7, SLC10A1, AFP, HP, HAMP, TTR, ONECUT1, SCTR

Schwann cells: SOX10, GAP43, S100B, NCAM1, NGFR

MAST-like, including basophils: SLC18A2, CPA3, MS4A2, GATA2, CMA1, CTSG, KIT, MRGPRX2, RGS13, MLPH, VWA5A, HPGDS, TPSAB1, TPSB2, SLC45A3, TPSG1

Myoblasts and myocytes: TTN, DLK1, NEB, DES, TNNT3, TNNI1, TCAP, TNNI2, KLHL41, MB, MYLPF, ACTC1, CKM, MYBPC2, CHRNA1, CADM2, MYF5

We examined the scores of these signatures on a range of original tumors examined manually, and found the following typical approximate values, set as expected values: TC 0.75, BC 0.6, MP 1.5, EP 1.3, CAF 1.2, EC 1, LIV 0.75, SC 1, MAST 1, MYO 0.5. We then performed auto-classification by scoring the signatures on all cells in datasets, dividing the scores by the expected values, and assigning cells to the category with the highest normalized score. This auto-classification was used to perform QC by cell type, which was done by setting a minimal and maximal mitochondrial gene content (typically from around 1 to 20%, but depends on the dataset and cell category) and number of detected genes (typically higher than 1-5 k, again depending on dataset and cell category). Definition of the thresholds was done by examination of scatter plots of mitochondrial content vs number of genes detected: the high quality live cells appear as a large cluster with high gene content and low mitochondrial content. A tail towards the top and/or left from this cluster would be dead and dying cells, and datapoints with low gene and/or mitochondrial content are debris, which were both excluded. We routinely re-adjusted the thresholds after closer examination of the cells during downstream analysis: well filtered cells cluster by cell type in dimensionality reduction rather than by quality, and failure to do so is typically seen as an indication that QC was not stringent enough and the analysis should be re-run.

We then re-imputed and scored the signatures on the filtered cells, and performed a refined manual cell-type classification by gating. This step naturally removes leftover debris and multiplets, which do not score high on any signature or score high on several signatures respectively. Of note, it is always possible that a cell category that was not considered might have been present in the dataset, and such cells would also score low on every category signature. We therefore examined the cells tagged for removal by looking for their markers and annotating the results in stringdb or searching for them in the literature before removing them. We considered datapoints with no unique markers, only combined markers of other cell types present in the dataset, to be debris/multiplets, whereas the presence of unique markers or of genes at their expression maximum on the suspected contaminants would warrant the addition of a new cell category.

We then computed a UMAP for the single dataset (not shown) for visualization.

When applicable (original tumors and genetically multiplexed co-cultures, or simple ID confirmation of mono-cultures for which WES is available), we looked for variants in the single-cell data, in particular as a way to distinguish cancer epithelium from non-mutated epithelium (marked “healthy” in the main figures). This was done with the burgertools custom functions, using the outputs obtained in the “Variant exploration at the single cell level” section. We opened the vcf file containing the variant info associated with the vartrix results with ReadVcf, which loads the results in a genotype object. We then added the VEP annotations to the genotype object with “AnnotateWithVEP”, and the TCGA prevalence of the mutations with AnnotateWithTCGA. We filtered variants for coverage >0 in at least 5 cells in the dataset with PrefilterVartrix, and perform a variant analogue of finding variable genes with the function FindInformativeVariants (which ranks variants based on their excess entropy, i.e. the entropy of the variant minus the trivial entropy expected from fraction of cells covered alone, which we found to be an excellent measure). We then examine the variants present in the dataset with the InformativeVariantPlot function (analogue to a VariableFeaturePlot), using the “summary” of the variants as annotations (i.e. Gene, position, impact), and filtering by impact (defined by VEP) and by prevalence in CRC according to TCGA to close-in on interesting mutations. We then examined how the data clustered based on genotypes rather than RNA expression with RunMDS (based on a fast divide-and-conquer implementation of MDS and a the custom genotype distance) and DimPlot, and examined the pattern of variant presence in both RNA and variant-based dimensionality reductions, with VariantPlot. We confirmed the identity of the patients when possible by comparison to the reference WES genotypes with CellSimilarityToGenotypes and VlnPlot. We then looked for the variants which were significantly different between the epithelial cells (or a subset of epithelial cells in rare cases when cancer cells were only a small fraction of the epithelial cells according to preliminary examination) and stromal cells, with FindDifferentVariantsTtest using a threshold of 20% difference in frequency and adjusted p-val < 0.05, and created a subset of the genotype object based on those, using the operator overloads for easy genotype handling. We used PseudoBulkGenotypes to compute aggregated genotypes by cell type, and examined the results with GenotypeSimilarityHeatmap to confirm the epithelial cells or a subset of epithelial cells did get distinguished from the stromal genotype. Then scored at the single cell level the similarity to the epithelial/cancer vs somatic genotype with CellSimilarityToGenotypes. In some cases, the results were too noisy (if only very few variants distinguish the cancer cells from others), and this could be overcome with CrossImpute, which uses the distances in a given assay (here RNA) to impute a quantity in another assay (here, the cell similarities to genotypes). We then gated the cancer vs healthy epithelial cells, as shown in Figure S4 panel C. We confirmed by manual examination that the classification was consistent with observed pathogenic mutations using the dimensionality reductions.

When applicable (datasets multiplexed with hashtag, i.e.. co-culture arrays), we performed demultiplexing based on hashtag oligos (HTOs). For this, we used again a gating approach packaged in burgertools. We normalized HTOs to their background level and expressed them as a percentage of the total HTOs per cell with NormalizeHTO, examined the cross-distribution of HTO expression with SignatureScatterPlot overlaid with cell types (which is informative as we knew which condition should contain which cell types in our co-culture experiments, which helps to fine-tune the tolerances), and applied manual gates with ClassifyManual. Cells with high percentage of several HTOs or not enough counts of any single HTO were thereby excluded.

Finally, we re-exported the filtered and annotated datasets for combined analysis downstream with the burgertools Export10X, which works in pair with Import10X for re-import. Of note, these functions export data from a Seurat object in the 10X format, i.e. mtx sparse matrix with barcodes and genes in associated tab separated values (tsv) files, compressed with gzip (gz), which is both efficient, and convenient for sharing and reuse across platforms and programming languages. Uniquely, these functions also export/import the metadata and dimensionality reductions as an additional metadata.tsv.gz file.

*Transcriptional network predictions*

We estimated transcriptional networks with the SCENIC workflow^8^, which includes the determination of modules based on co-expression and their TF regulators based on the presence of cis-regulatory motifs.

We re-exported a balanced subset of epithelial cells from all OTs and PDOs for regulon analysis. We imputed the full combined epithelial portions of the datasets with MAGIC. We then set the genes with an expression value of less than 0.02 to 0 in order to reform a sparse matrix for computation efficiency and because very low values have no meaning. We then subset the genes to keep only the 8 k most variable on this subset (which we found to be the ones truly showing contrast between populations rather than purely stochastic), and kept a maximum of 100 healthy and 100 cancer cells per dataset for each of the 63 datasets. This has the double interest of making the calculations tractable, and of balancing the influence of various datasets, and of cancer vs healthy epithelium, which we found to lead to the most informative predicted networks due to the well represented cell diversity.

We then computed the predicted transcription networks with the SCENIC functions pyscenic grn and pyscenic ctx, using the dabases in their version 10 (hg38_10kbp_up_10kbp_down_full_tx_v10 and hg38_500bp_up_100bp_down_full_tx_v10). We picked up the results in R, and scored the regulons with both AUC and burgertools. We found the time of computation with AUC to be a major hurdle, and the simpler scores from burgertools to be equally informative, and therefore used the latter. We computed the variance of the scores from *in vivo* vs *in vitro*, patients, and cancer vs healthy epithelium, and plotted the subnetworks obtained with various filters (such as a target gene, the directionality of the regulation, or applying a max number of target genes per transcription factor to exclude near-ubiquitous regulators) with igraph and visNetwork. An interactive explorer where the filters can be adjusted by users is available on the CRC-TME online atlas.

*Cell-cell interaction predictions*

We predicted cell-cell interactions for each OT dataset individually with CellPhoneDB v4. We then combined the results across patients, and appended metadata such as gene expression by cell type, patients for which the interaction exists, CMS of these patients etc with custom R code, that also reformatted the results for visualization as a network, filtering by cell types with custom thresholds, and export to cytoscape for fine-tuning of the network layout (i.e. force directed layout, with manual adjustments to maximize visibility, and annotations overlaid in Illustrator). We also provide an interactive explorer of this network in which users can adjust the custom filters on the CRC-TME online atlas. The app, written in javascript and relying on cytoscape.js, lets users quickly narrow down on any interaction for which the ligand or receptor are expressed (or not) on cell types of interest, or select subnetworks/neighbors, close-up, subset, re-arrange manually etc.

*CMS classification*

We predicted the CMS of OTs in R with the library CMSclassifier (https://github.com/Sage-Bionetworks/CMSclassifier), based on pseudo-bulk aggregation of the scRNA-seq data. We evaluated the contribution of single cell types to the overall score by performing the same scoring at the single cell level. We estimated the intrinsic CMS (iCMS) by performing the pseudo-bulk aggregation on epithelial cells only.

*Enriched pathways in OT vs mono-cultures and by celltype between culture conditions in co-cultures*

We computed enriched pathways between OTs and mono-cultures in R using the log2FC as a metric, with the library fgsea. We selected a subset of MSigDB Hallmarks, KEGG pathways, and Reactome pathways by excluding pathways which were entirely irrelevant (e.g. related to other diseases), and then removing duplicates which cover the same pathway across different databases, taking in order of priority the pathways from Hallmarks>KEGG>Reactome, which led us to a list of 433 unique pathways. The top results were shown in heatmaps created with the R library ComplexHeatmap 2.14.0.

*Differential expression analysis*

Differential expression between various populations were computed with Seurat FindMarkers, using log in base 2 and a pseudo-count of 0.01 (data normalized to 10 k), for individual cell types between two different conditions within single datasets (in co-cultures) or for a given cell type between two datasets (OT vs monocultures), and averaged step by step as indicated in figure legends to generate summarized data (e.g. across replicates, then across patients). Each cell was given the same weight when averaging across replicates, each patient was given the same weight when averaging across patients. Individual plots without averaging, and manual adjustment of the filtering thresholds can be done on the CRC-TME online atlas. In co-culture experiments, genes in a comparison were considered significant for a given patient if they were significant in at least two of the replicates. Averaged volcano plots were created with ggplot2 v3.4.1 by plotting the significance *vs*. Log2 fold change, and LFC-LFC plots by plotting two Log2 fold changes from different effects against each other.

*Comparison of PDO response to individual factors vs co-cultures*

In order to compare the response of PDOs to individual factors to their response to a given cell type in co-cultures, we pre-grew PDOs from 10 k cells in 20 µl Matrigel domes in 24 well plates until they reach a size of a few hundred µm (6 to 12 days depending on the patient), then exposed them to the individual factors listed in Figure 7E for 3 days. We then collected the cell-laden hydrogels in Trizol and proceeded to RNA isolation, followed by bulk RNA-seq with BRB-seq^9^ and Illumina sequencing (HiSeq). We obtained 720 k usable reads per sample on average. We repeated the screen for 3 patients. The data was aligned to GRCh38 with star 2.7.6a. We then compared the log2 fold change due to exposure to an individual factor (comparing to a control condition with no treatment, done in quadruplicate) to that due to exposure to a co-culture with one or several other cell types. We fitted the LFC-LFC plot with a linear model in R, restricting the curve fitting to genes which were significantly differentially expressed in at least one patient for the response to the cell type in question in the co-culture experiments (genes highlighted in an example in Figure 7D), to limit the noise from the mass of low-expressed genes and focus the analysis on the genes of interest. The coefficient of linearity and coefficient of determination (R^2^) from these fits were used to construct a dot plot with ggplot2, highlighting which individual factors could mimic the influence of a given co-culture.

*T cell activity monitoring with cytometric bead array*

In order to monitor T cell inhibition by CAFs, we seeded 10 k CAFs and 25 k TILs per well in round-bottom low-adhesion 384 well plates, in 50 µl of MV:hCoBM 1:1 + IL2 + CD3/28 mAbs. Controls included conditions with no CAFs or no activation by mAbs, or replacing the CAFs by PDO fragments deposited on a 5 µl layer of Matrigel (flattened by centrifugation). We analyzed the supernatant at D2 and D5, quantifying the concentrations of cytokines that reflect T cell activation, namely IFNG, TNF, and Granzyme B, using a cytometric bead array readout (CBA, BD Bioscience 560111, 558273, 560304). We analyzed the results with a multivariate ANOVA, considering the combined measurements on each sample (i.e. 6 scalar values for 2 days and 3 cytokines) as one datapoint, with a Pillai post-hoc test and false discovery rate control, comparing all conditions condition to the activated positive control. The non-activated T cells served as an additional control of T cell functionality and responsiveness at the time of the experiment.

*References*

1.	Rezakhani, S., Gjorevski, N., Lutolf, M.P., Rezakhani, S., Gjorevski, N., and Lutolf, M.P. (2020). Low-Defect Thiol-Michael Addition Hydrogels as Matrigel Substitutes for Epithelial Organoid Derivation. Adv Funct Mater *30*, 2000761. 10.1002/ADFM.202000761.
